# Resistance to Radiation Enhances Metastasis by Altering RNA Metabolism

**DOI:** 10.1101/2025.02.19.638943

**Authors:** Ayush Kumar, Kensei Kishimoto, Hira Lal Goel, Christi A. Wisniewski, Rui Li, Brendan Pacheco, Lihua Julie Zhu, William A. Flavahan, Arthur M Mercurio

**Affiliations:** Departments of Molecular, Cell and Cancer Biology, University of Massachusetts Chan Medical School, Worcester MA

## Abstract

The cellular programs that mediate therapy resistance are often important drivers of metastasis, a phenomenon that needs to be understood better to improve screening and treatment options for cancer patients. Although this issue has been studied extensively for chemotherapy, less is known about a causal link between resistance to radiation therapy and metastasis. We investigated this problem in triple-negative breast cancer (TNBC) and established that radiation resistant tumor cells have enhanced metastatic capacity, especially to bone. Resistance to radiation increases the expression of integrin β3 (ITGβ3), which promotes enhanced migration and invasion. Bioinformatic analysis and subsequent experimentation revealed an enrichment of RNA metabolism pathways that stabilize ITGβ3 transcripts. Specifically, the RNA binding protein heterogenous nuclear ribonucleoprotein L (HNRNPL), whose expression is regulated by Nrf2, mediates the formation of circular RNAs (circRNAs) that function as competing endogenous RNAs (ceRNAs) for the family of let-7 microRNAs that target ITGβ3. Collectively, our findings identify a novel mechanism of radiation-induced metastasis that is driven by alterations in RNA metabolism.

## Introduction

Metastasis is a multifactorial process that requires functional reprogramming in tumor cells that enable them to migrate, survive in circulation and colonize distal sites. Key molecular pathways that tumor cells utilize in each individual stage of the metastatic process have been identified, but how these pathways are executed is less clear. There is evidence that the process is stochastic and a result of intratumor heterogeneity (1, 2), as well as data that it involves sequential adaptations to extrinsic factors such as biomechanical forces (3, 4), the tumor microenvironment (5, 6), and treatment exposure (7, 8). We are particularly interested in the role of treatment in enhancing the metastatic process because advances in multi-omics technology and cellular tracking at a single-cell level have demonstrated a significant overlap between therapy resistance and metastasis, especially in the context of chemotherapy (9, 10). In other terms, understanding the role of therapeutic modalities in selecting for the survival of resistant cells that have a greater propensity to metastasize is a challenging biological problem with profound consequences for patient outcome.

The link between resistance to radiation therapy and metastasis has not been studied extensively (11–13). This issue is particularly relevant for triple-negative breast cancer (TNBC) because radiation is a reliable therapy option because it can reduce locoregional recurrence and decrease mortality rates (14, 15). However, there is a subset of patients for whom adjuvant radiotherapy increases the number of circulating tumor cells and is associated with an increased likelihood of distant metastasis (16). Thus, it is critical to explore the mechanisms involved in resistance to radiotherapy in TNBC and their contribution to distant metastasis. To address this issue, we developed radioresistant models of TNBC and demonstrated their enhanced metastatic potential *in vivo*. The data we obtained using these models reveal that changes in RNA metabolism, especially the regulation of circular RNAs, underlie radiation resistance and the propensity to metastasize because they enable the expression of a key effector of these processes, the integrin β3 (ITGβ3).

## Results

### Radiation resistance enhances metastasis

To investigate resistance to radiation in a consistent and reproducible manner, we utilized two TNBC cell lines, one mouse (4T1) and one human (MDA-MB-468), and implemented a dose escalation radiation strategy (**Fig S1A**). This approach has been utilized in other studies to investigate resistance to radiation and delivers a biologically equivalent dose of radiation given to patients (17). The irradiated cell lines (MDA-MB-468-RR and 4T1-RR) were radioresistant compared to the parental cell lines as evidenced by the higher surviving fraction in a clonogenic assay and reduced apoptosis seen at 8Gy based on Annexin V staining (**Fig 1A**, **1B Fig S1B**). To understand mechanisms of resistance within these cells, we performed transcriptomic analysis. These studies showed that radioresistant cells were quite different from the parental cells in the 4T1 system; however, there were fewer changes in the MDA-MB-468 system despite them both receiving the same radiation dose and developing a radioresistant phenotype (**Fig S1C**). Interestingly, gene set enrichment analysis indicated an upregulation of several curated metastasis gene sets in the radioresistant (RR) cell lines scompared to the parental cell lines (**Fig 1C**). Using Transwell invasion and scratch-wound migration assays, we found that the radioresistant cells had an enhanced invasive and migratory capacity, properties associated with metastatic cells. (**Fig 1D-F**).

**Figure 1:**
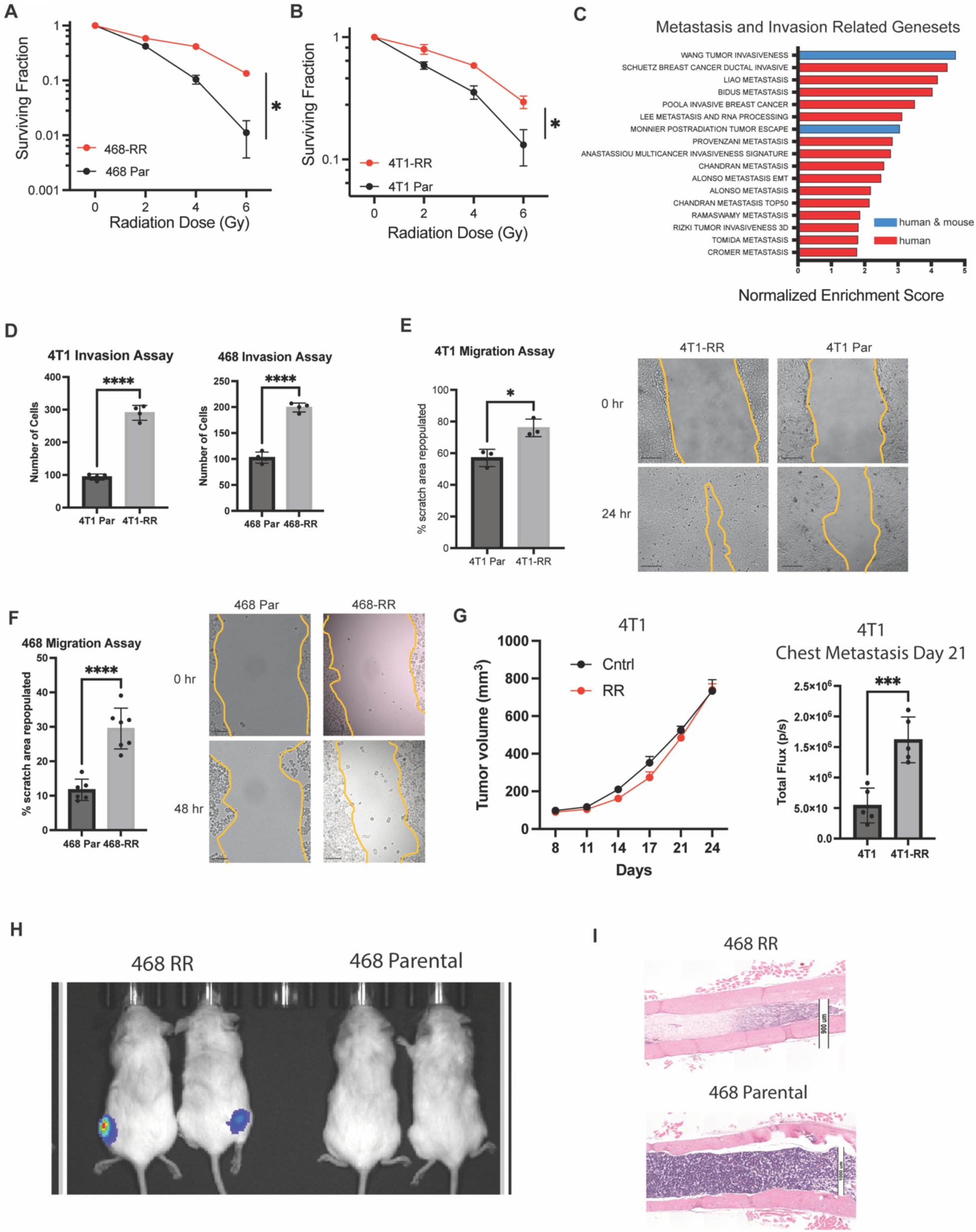
Radiation resistance enhances metastasis. **A.** A clonogenic assay of MDA-MB-468-RR vs MDA-MB-468 showing the surviving fraction after radiation of 0, 2, 4, and 6Gy (n=3 biological replicates, representative experiment, mean ± SD). **B.** A clonogenic assay of 4T1-RR vs 4T1 showing the surviving fraction after radiation of 0, 2, 4, and 6Gy (n=3 biological replicates, representative experiment, mean ± SD). **C.** The normalized enrichment score for genesets associated with metastasis, invasion, and migration that were upregulated in the radioresistant cell lines compared to the parental cell lines (blue is found in both human and mouse genesets whereas red is found in the human genesets). **D.** The number of cells that were able to invade through the Matrigel of the Transwell in the radioresistant and parental cell lines (n=4, mean± SD, student’s t-test). **E.** The percentage of the scratch area that was repopulated after 24 hours for the 4T1 cells (n=3, mean± SD, student’s t-test, scale bar is 200um) and **F.** 48 hours for the MDA-MB-468 cells (n=3, mean± SD, student’s t-test, scale bar is 200um). **G.** The tumor growth of the 4T1 and 4T1-RR tumors in the mammary fat pad of Balb/c mice after orthotopic injection and the total flux in the thoracic region of the mice that was seen 21 days after implantation (n=5, mean± SEM, ANOVA 2-way). **H.** The bioluminescent image taken 64 days after intracardiac injection of MDA-MB-468-RR and MDA-MB-468 cells (n=2, 2×10^5^ cells/injection). **I.** Histological images of the femur taken from the mice. The scale bar is 1000um for the upper image and 900um for the lower image.

Next, we compared the metastatic potential of RR cells to their parental controls. Orthotopic injection of 4T1-RR in the mammary fat pad of BALB/C mice resulted in increased chest metastasis compared to the 4T1 cells, despite having no difference in primary tumor growth (**Fig 1G**). Histological analysis of these mice revealed more macro-metastatic sites in the thoracic cavity of mice with 4T1-RR tumors relative to the ones with 4T1 tumors (**Fig S1D**). To explore hematogenous metastasis, mice were given intracardiac injections of either MDA-MB-468-RR or MDA-MB-468 cells to observe multi-organ spread. One of the salient differences observed was the presence of bone metastasis with MDA-MB-468-RR cells and not with MDA-MB-468 cells as evidenced by bioluminescent imaging and histological analysis (**Fig 1H-I, S1E**). These findings support the hypothesis that resistance to radiation is associated with enhanced metastasis, especially hematogenous metastasis to bone.

### Unbiased identification of integrin β3 as a driver of radiation-induced metastasis

To uncover the mechanism by which radiation resistance enhances metastasis, we curated the upregulated genes that were expressed in both the MDA-MB-468-RR and 4T1-RR cell lines (**Fig 2A**). Gene ontology analysis of these shared genes revealed blood coagulation and integrin signaling as enriched terms with ITGβ3 (integrin β3) as a gene common to both (**Fig 2B**). This finding is consistent with previous work implicating ITGβ3 as an important modulator of radiotherapy resistance (18, 19) and metastasis (20, 21). To verify our bioinformatic data, we observed a significant increase in ITGβ3 mRNA, protein, and cell surface expression (**Fig 2C-D, S2A**) in the MDA-MB-468-RR compared to MDA-MB-468 cells. Similar results were observed comparing 4T1 and 4T1-RR cells (**Fig 2E-F, S2B**). Pretreating the 468-RR cells with an integrin β3 function blocking antibody decreased the migration capacity of the cells (**Fig 2G**) and diminishing ITGβ3 expression of 4T1-RR cells using RNA interference decreased the migration of 4T1-RR cells (**FigS2C**).

**Figure 2.**
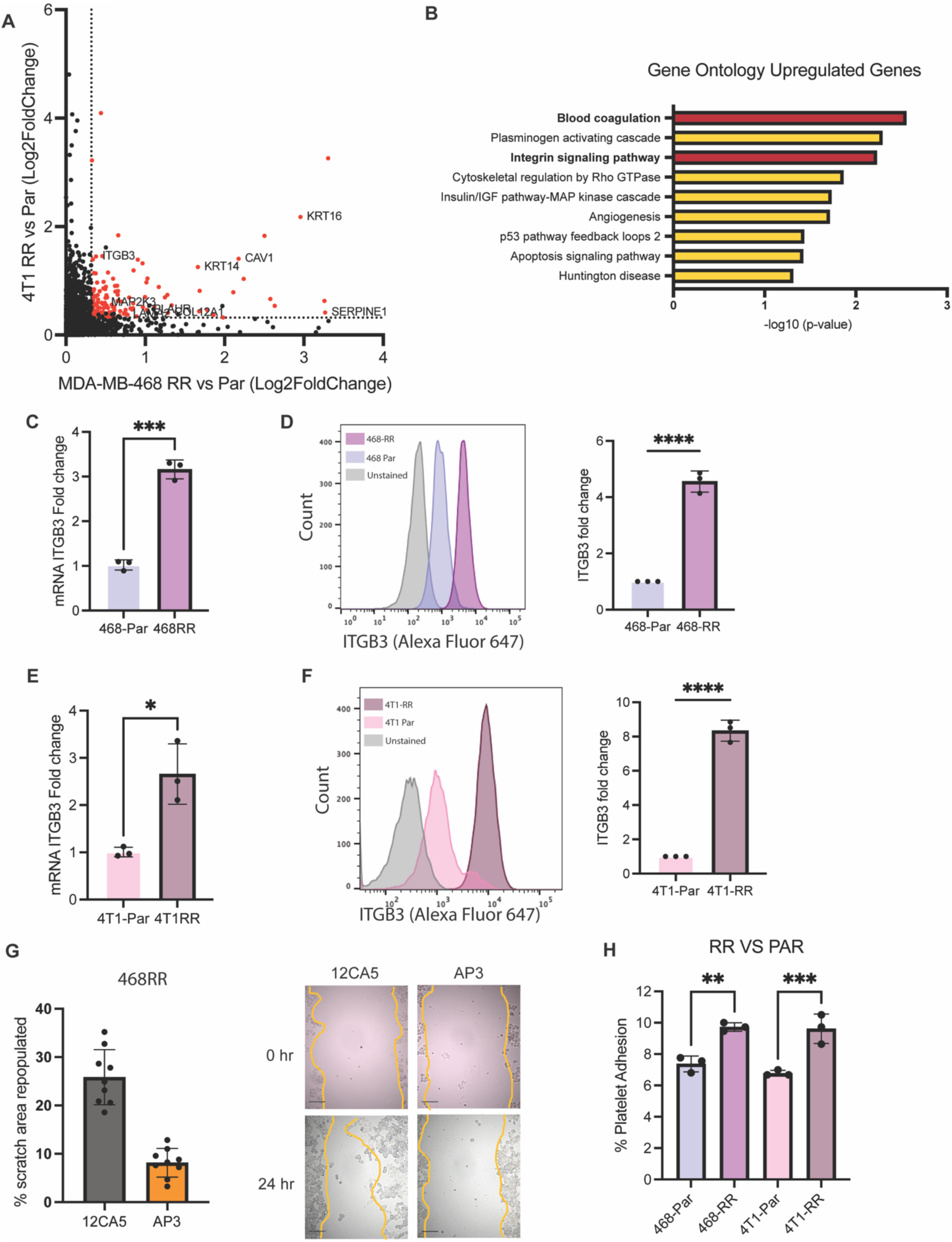
Unbiased identification of integrin β3 as a driver of radiation-induced migration and invasion. **A.** A graph of the differentially upregulated genes based on the Log2 fold change when comparing the radioresistant cell lines to the parental cell lines. The red dots are the 135 upregulated genes in both cell lines that meet the criteria of a FC>1.25 and FDR<0.1 **B.** The gene ontology analysis of the 135 shared genes identified in Fig 2A with ‘blood coagulation’ and ‘Integrin signaling pathway’ being enriched. **C.** The mRNA expression of ITGβ3 based on RT-qPCR (n=3, mean ± SD, unpaired t-test two-tailed) and **D.** the surface protein expression of ITGβ3 based on flow cytometry in the MDA-MB-468-RR and MDA-MB-468 cells (n=3, mean ± SD, unpaired t-test two-tailed). **E.** The mRNA expression of ITGβ3 based on RT-qPCR (n=3, mean ± SD, unpaired t-test two-tailed) and **F.** the surface protein expression of ITGβ3 based on flow cytometry in the 4T1-RR and 4T1 cells (n=3, mean ± SD, unpaired t-test two-tailed). **G.** The percentage of the scratch area that was repopulated after 48 hours for the MDA-MB-468 cell lines after treating with an integrin β3 function blocking antibody (AP3) and its appropriate control (12CA5) (n=9, mean ± SD, unpaired t-test two-tailed). The scale bar is 200um. **H.** The percent of platelets that were able to bind to the MDA-MB-468 and 4T1 cell lines seen *in vitro*. n=3, mean ± SD, and unpaired t-test two-tailed.

Given that binding of platelets to tumor cells can facilitate metastasis (22–24) and that ITGβ3 was associated with the gene ontology term of ‘blood coagulation’, we investigated the possibility that this integrin promotes the binding of RR cells to platelets. Indeed, we observed that the RR cell line had an increased capacity to bind to human platelets *in vitro* (**Fig2H**). These data demonstrate the necessity of ITGβ3 for the migration and metastasis of radioresistant cells. **HNRNPL stabilizes ITGβ3 mRNA:** Further analysis of the gene sets that are enriched in the RR cell lines revealed an intriguing pattern of RNA stability and metabolism (**Fig 3A, S3A**), suggesting that mRNA stability may contribute to ITGβ3 expression and, consequently, metastasis. To investigate this hypothesis, we treated cells with actinomycin D and then quantified ITGβ3 mRNA. We observed that the RR cells had a higher level of ITGβ3 compared to the parental cells when transcription was inhibited with actinomycin D (**Fig S3B**). To identify potential regulators of ITGβ3 mRNA stability, we identified four genes from the RNA stability gene set whose expression was enhanced in both of the radioresistant cell lines compared to their controls, one of which was heterogeneous ribonucleoprotein L (HNRNPL). The transcript levels of HNRPNL are strongly correlated with ITGβ3 mRNA expression and are a strong predictor of overall survival in breast cancer (**Fig S3C**); however, their role in the context of radiation resistance and metastasis has not been investigated. Indeed, we observed a significant increase in HNRNPL mRNA and protein expression in both MDA-MB-468-RR and 4T1-RR cells compared to their controls (**Fig 3B and 3C**). Also, a strong positive correlation between HNRNPL and ITGβ3 expression was seen in breast cancer patients (**Fig S3C**).

**Figure 3.**
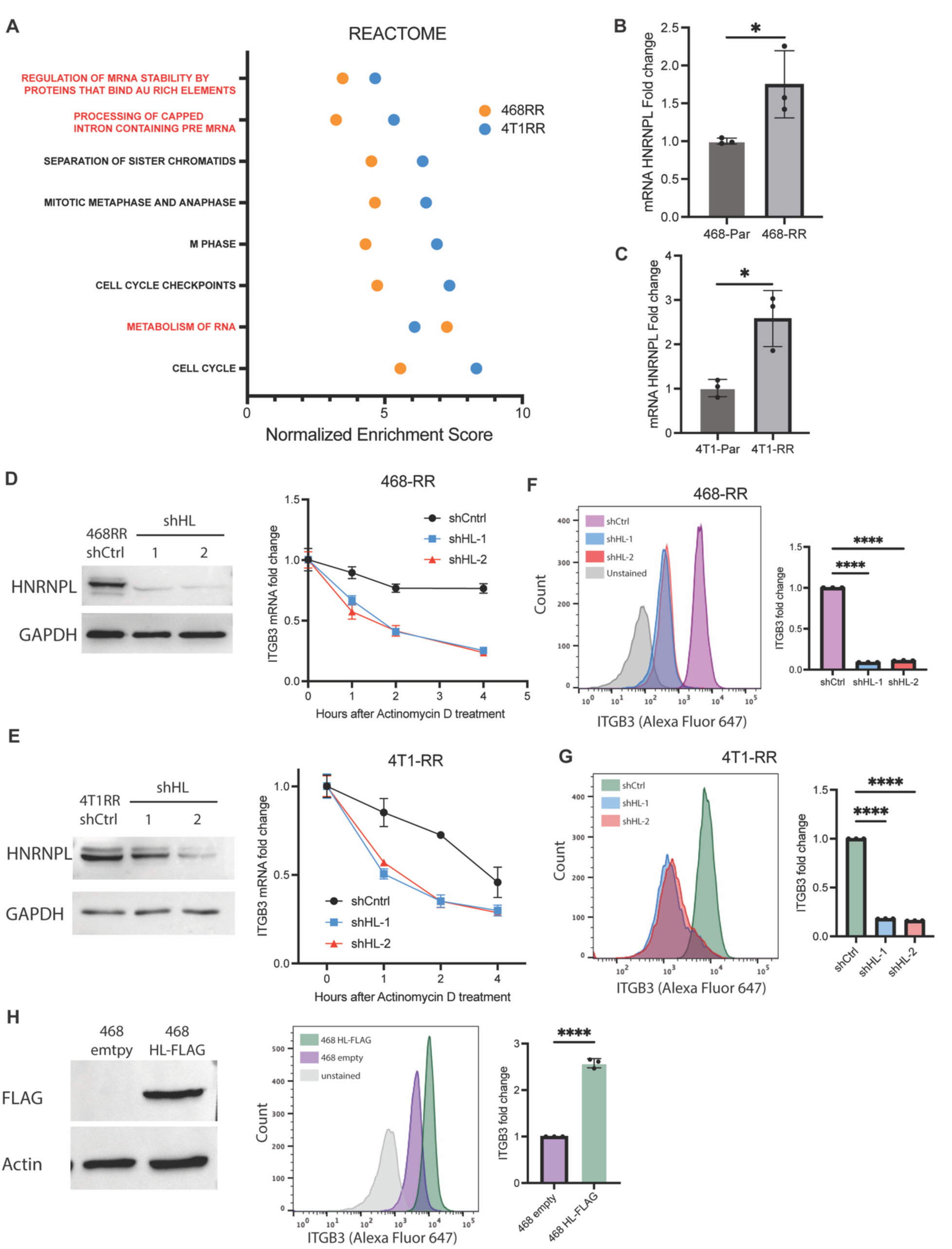
HNRNPL stabilizes ITGβ3 mRNA and regulates integrin β3 expression. **A.** The gene sets from ‘REACTOME’ that were upregulated in the 4T1-RR and MDA-MB-468-RR cells with the red signifying gene sets associated with RNA metabolism. The mRNA expression of HNRNPL based on RT-qPCR in the **B.** MDA-MB-468 cell lines (n=3, mean ± SD, unpaired t-test two-tailed) and the **C.** 4T1 cell lines (n=3, mean ± SD, unpaired t-test two-tailed). **D.** The protein expression of HNRNPL in MDA-MB-468-RR cells and the HNRNPL-knockdown cells (shHL-1 and shHL-2) and the transcript stability assay of ITGβ3 after 0,1,2, and 4 hours of actinomycin D treatment (n=3, mean ± SD, two-way ANOVA with Tukey’s multiple comparison tests). **E.** The protein expression of HNRNPL in 4T1-RR cells and the HNRNPL-knockdown cells (shHL-1 and shHL-2) and the transcript stability assay of ITGβ3 after 0,1,2, and 4 hours of actinomycin D treatment (n=3, mean ± SD, two-way ANOVA with Tukey’s multiple comparison tests). **F.** The surface expression of ITGβ3 in the MDA-MB-468-RR cells and the HNRNPL-knockdown cells (n=3, mean ± SD, one-way ANOVA with Dunnett’s multiple comparisons tests). **G.** The surface expression of ITGβ3 in the 4T1-RR cells and the HNRNPL-knockdown cells (n=3, mean ± SD, one-way ANOVA with Dunnett’s multiple comparisons tests). **H.** The western blot expression of FLAG in the MDA-MB-468 cells transfected with empty vector and FLAG-tagged HNRNPL. The surface expression of ITGβ3 in the MDA-MB-468 empty vector and HNRNPL overexpression cells (n=3, mean ± SD, unpaired t-test two-tailed).

To investigate whether HNRNPL mediates integrin β3 expression, we used RNA interference to reduce HNRNPL expression and then assessed ITGβ3 mRNA stability. A significant decrease in ITGβ3 transcripts after actinomycin D treatment was observed in RR cells in which HNRNPL expression had been diminished compared to control cells (**Fig 3D and 3E**). Moreover, diminishing HNRNPL expression also decreased integrin β3 protein and surface expression (**Fig 3F-G, S3D-E**). The expression of HNRNPL was diminished in the parental MDA-MB-468 cells and 4T1 cells, which also demonstrated a decrease in integrin β3 surface expression (**Fig S3F-G**). Increasing the expression of HNRNPL in T47D and MDA-MB-468 cells resulted in a significant increase in the surface expression of integrin β3 (**Fig 3H, S3H-I**). The data implicate HNRNPL as key mediator of ITGβ3 mRNA stability and, consequently, protein surface expression in radioresistant TNBC.

### NRF2 regulates HNRNPL transcription

Next, we sought to identify the upstream factors responsible for increased HNRNPL expression in response to radiation resistance. Our RNA-seq analysis enabled us to identify 289 transcription factors that were differentially active in RR compared to control cells. We then used ChIP-seq data from ENCODE to identify transcription factors that bind near the promoter region of HNRNPL. This screening strategy identified 17 possible transcription factors in RR cells that could regulate HNRNPL expression (**Fig 4A**). Among these 24 factors, we noted that the NRF2/KEAP1 pathway was enriched in the RR cells (**Fig S4A**). Moreover, there is a positive correlation between NRF2 mRNA expression and HNRNPL mRNA expression in basal breast cancer patients (**Fig S4B**). Given that multiple regulatory components of NRF2 activation are independent of its transcript levels (25), we utilized TCGA data to generate a NRF2 activity score, based on a 16-gene signature (26), which had a positive correlation with HNRNPL mRNA expression (**Fig 4B**). From ChIP-seq data from ENCODE, we identified a NRF2 binding site near the HNRNPL promoter (**Fig S4C**). We compared NRF2 activity differences between the MDA-MB-469-RR and control MDA-MB-468 cells based on the ratio of nuclear to cytoplasmic localization of NRF2 and observed a significant increase in NRF2 activity in the radioresistant cells compared to the parental 468 cells (**Fig 4C**). We diminished the expression of NRF2 and observed a decrease in HNRNPL expression at both the mRNA levels in MDA-MB-468-RR (**Fig 4D**). The NRF2 knockdown 4T1-RR cells also had decreased expression of HNRNPL (**Fig 4E**). We utilized a NRF2 inhibitor, ML385 (27), which demonstrated a decrease in HNRNPL expression after 72 hours of treatment in the MDA-MB-468-RR cells compared to the DMSO-treated control (**Fig 4F**). Subsequently, we treated parental MDA-MB-468 cells with dimethyl fumarate (DMF), which has been shown to induce NRF2 activity after 24 hours (28). We observed an increased expression of the Nrf2 target gene HMOX1 and an associated increase in HNRNPL transcripts (**Fig 4G**). After establishing the role of NRF2 in regulating HNRNPL, we used ChIP-qPCR which provided evidence that NRF2 binds to the promoter region of HNRNPL that is identified in Figure S4B (**Fig 4H**). Thus, the data show that the enhanced activity of NRF2 in the radioresistant cells provides a migratory advantage in the cells via HNRNPL expression.

**Figure 4.**
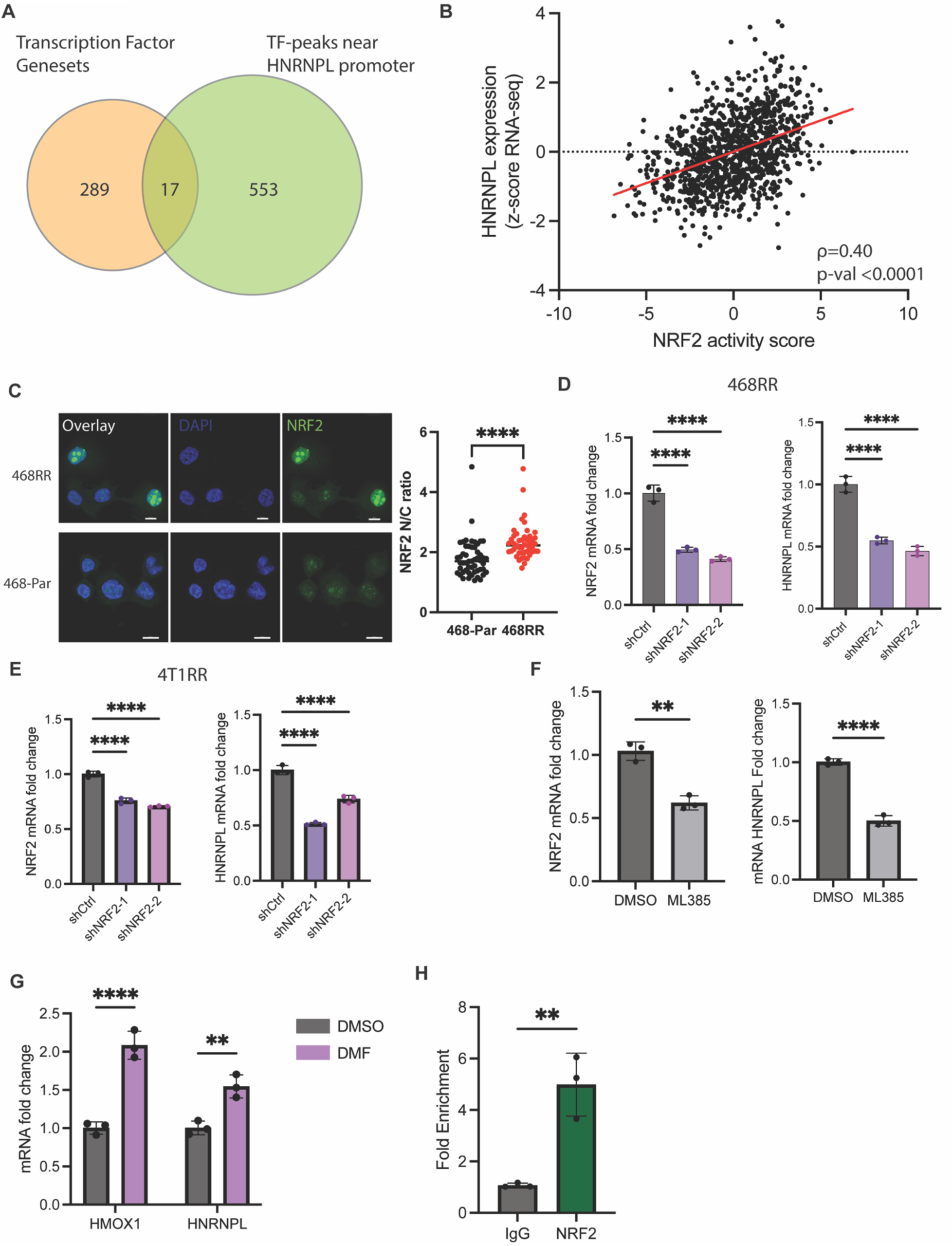
NRF2 regulates HNRNPL transcription. **A.** A Venn diagram showing the transcription factor genesets that were upregulated based on GSEA of the RNA-seq comparing the radioresistant cell lines to the parental cell lines along with the transcription factors that were identified by ENCODE transcription factor ChIP-seq data to regulate HNRNPL. B. A correlation between NRF2 activity score compared to HNRNPL expression in breast cancer patient samples collected by TCGA (n=1108, Spearman correlation). **C.** Immunofluorescent images of the MDA-MB-468-RR and MDA-MB-468 cells with blue being the DAPI dye and green representing NRF2. Scale bar is 10um. A ratio comparing NRF2 localization in the nucleus vs the cytoplasm for the MDA-MB-468-RR (n=52) vs MDA-MB-468 cells (n=47). Data represented as mean ± SD, and statistical analysis involved one-way ANOVA with Dunnett’s multiple comparisons tests. **C.** The mRNA expression of NRF2 and HNRNPL for the MDA-MB-468-RR and NRF2-knockdown cells (shNRF2-1 and shNRF2-2). n=3, mean ± SD, one-way ANOVA with Dunnett’s multiple comparisons tests. **D.** The mRNA expression of NRF2 and HNRNPL for the 4T1-RR and NRF2-knockdown cells (shNRF2-1 and shNRF2-2). n=3, mean ± SD, one-way ANOVA with Dunnett’s multiple comparisons tests. **E.** The mRNA expression of NRF2 and HNRNPL for the MDA-MB-468-RR cells treated with DMSO or ML385 (5uM) for 72 hours (n=3, mean ± SD, one-way ANOVA with Dunnett’s multiple comparisons tests). **F.** The mRNA expression of NRF2 and HNRNPL for the MDA-MB-468 cells treated with DMSO or DMF (5uM) for 24 hours (n=3, mean ± SD, unpaired t-test two tailed). **G.** The fold enrichment of NRF2 binding to the HNRNPL promoter region identified by ChIP-qPCR when pulling down NRF2 vs IgG (n=3, mean ± SD, unpaired t-test two tailed).

### HNRNPL-mediated circular RNA formation regulates ITGβ3

HNRNPL can impact RNA processing by alternative splicing of mRNAs and by forming circular RNAs (circRNAs). To assess alternative splicing, we examined publicly available CLIP-seq databases of HNRNPL (29), as well as an RNA binding protein prediction algorithm, RBPDB (30), to identify potential ITGβ3 binding sites (**Fig S4D**). We performed RIP-qPCR for RNA bound to HNRNPL in MDA-MB-468-RR cells using the representative peaks, but we could not identify a direct interaction between HNRNPL and the ITGβ3 transcript (**Fig S4E**). Therefore, we focused our efforts on identifying the circRNAs whose expression is mediated by HNRNPL in MDA-MB-468-RR cells. CircRNAs formed by HNRNPL can function as competing endogenous RNAs (ceRNAs) that can scavenge miRNAs (31). To identify such ceRNAs, we used Induro-RT mediated circRNA sequencing (IMCR-seq), a new strategy to identify different isoforms of circRNAs (32) that are expressed in the MDA-MB-468-RR cells and the HNRNPL knockdown cells and had high correlation with biological samples (**Fig S5A**). Several differentially expressed circRNAs were identified, and we focused on identifying the circular RNAs upregulated in the radioresistant cells (**Fig 5A**). To assess their ability to scavenge miRNAs that target ITGβ3, we used Circr, a computational tool for the prediction of miRNA:circRNA interactions (33), and identified binding sites for let-7 miRNAs for the top differentially expressed ceRNAs. We focused our analysis on let-7 miRNAs because of their conservation across species (34), role in breast cancer tumorigenicity (35), and previous studies had elaborated their role in the regulation of ITGβ3 transcript stability (36, 37). To screen for the important ceRNA regulators, we developed a score for each circRNA isoform that takes into the account the number of let-7 binding sites, its differential expression between the HNRNPL knockdown cells and the MDA-MB-468-RR cells, and its abundance in the radioresistant cell line (**Fig 5B, Supplementary Table 1**). From this score, we identified three circular RNAs that are highly expressed in MDA-MB-468-RR compared to HNRNPL knockdown cells that we validated by RT-qPCR: *circRAB12*(*2,3,4,5,6*), *circBACE2(2,3,4,5,6,7,8)*, and *circBIRC6(2,3,4,5,6)*. (**Fig 5C**). Furthermore, we found that increasing the expression of HNRNPL in MDA-MB-468 and T47D cells increases the expression of these circRNAs (**Fig 5D, S5B**). By diminishing the expression of the ceRNAs with siRNAs, we achieved a significant decrease in ITGβ3 mRNA and protein expression (**Fig 5E, S5C**). Lastly, using a lentiviral plasmid that generates scarless circular RNAs (38) to express *circRAB12*, which is one of the ceRNAs predicted to bind to let-7 microRNAs, in the HNRNPL knockdown cells, we observed a significant increase in ITGβ3 mRNA and surface expression of β3 compared to the empty plasmid control (**Fig 5F, S5D**). Taken together, these data provide significant evidence that the HNRNPL-derived circular RNAs in radiation resistant cell lines have the capacity to sponge microRNAs that target ITGβ3, and consequently, enhance integrin β3 expression.

**Figure 5.**
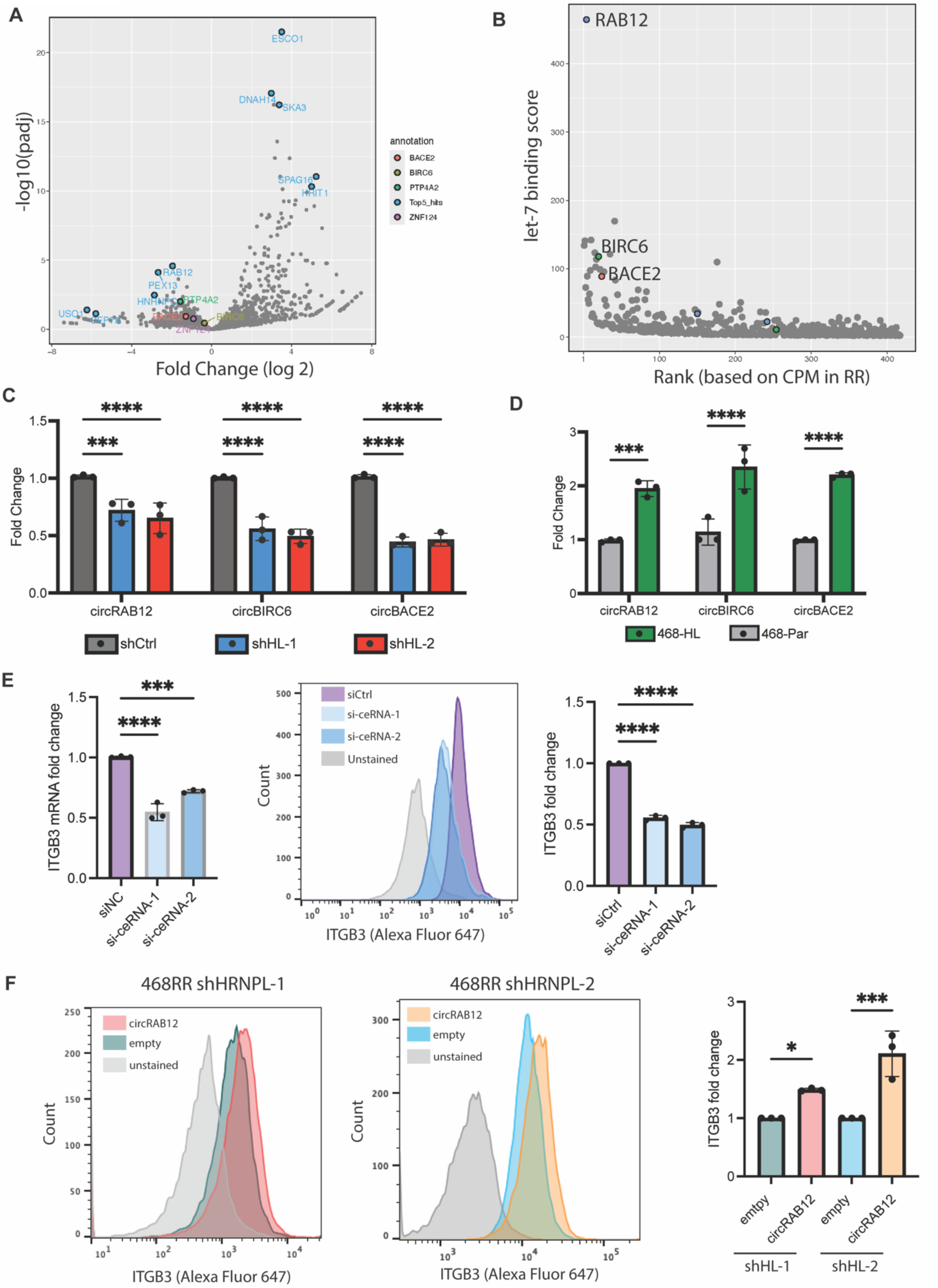
HNRNPL-mediated circular RNA formation regulates ITGβ3. **A.** IMCR-seq results showing the differential expression of different circular RNAs in the MDA-MB-468-RR and in the shHNRNPL-1 cells. The blue dots highlight the top 5 differentially expressed circular RNAs whereas the other labels represent other circular RNAs that can function as ceRNAs **B.** A graph showing the let-7 binding score for each circular RNA in comparison to its abundance (based on ranking of circular RNAs from CPM values) in MDA-MB-468-RR cells. **C.** mRNA expression of the top ceRNAs identified by Circr in the MDA-MB-468-RR and HNRNPL-knockdown cells (n=3, mean ± SD, one-way ANOVA with Dunnett’s multiple comparisons tests). **D.** mRNA expression of the top ceRNAs in MDA-MB-468 transfected with empty vector and HNRNPL (n=3, mean ± SD, unpaired t-test two-tailed). **E.** mRNA expression of ITGβ3 after siRNA treatment of the ceRNAs, with a representative image of the surface expression of integrin β3 and quantification based on flow cytometry (n=3, mean ± SD, unpaired t-test two-tailed). **F.** Representative flow cytometry data showing the surface expression of integrin β3 in the HNRNPL knockdown cells transducing with an empty plasmid or *circRAB12*. The graphs quantify the fold change in integrin β3 surface expression in the HNRNPL knockdown cells after transducing them with an empty plasmid or *circRAB12.* n=3, mean ± SD, unpaired t-test two-tailed.

### HNRNPL-derived circRNAs drive migration by stabilizing ITGβ3 mRNA

The previous data provided a strong rationale for HNRNPL-mediated stability of ITGβ3 transcripts, and we wanted to observe if it contributed to migration in radioresistant cells. The HNRNPL knockdown cells were less migratory compared to the radioresistant cells (**Fig 6A and 6B**). These HNRNPL knockdown cells were able to rescue the migratory capacity by overexpressing ITGβ3 with a plasmid (**Fig 6C and 6D**). Moreover, we decreased expression of the ceRNAs that regulate ITGβ3 in MDA-MB-468-RR cells and observed a decreased migration capacity compared to the control (**Fig 6E**). To further elucidate the role of HNRNPL in mediating metastasis via ITGβ3, we explored its capacity to regulate tumor-platelet clusters. We observed a significant decrease in attached platelets to the HNRNPL knockdown cells compared to the radioresistant cells (**Fig 6F**). Overall, we were able to characterize the functional capacity of radioresistant cells to migrate and attach to platelets via the HNRNPL/circRNA/ITGβ3 regulation.

**Figure 6.**
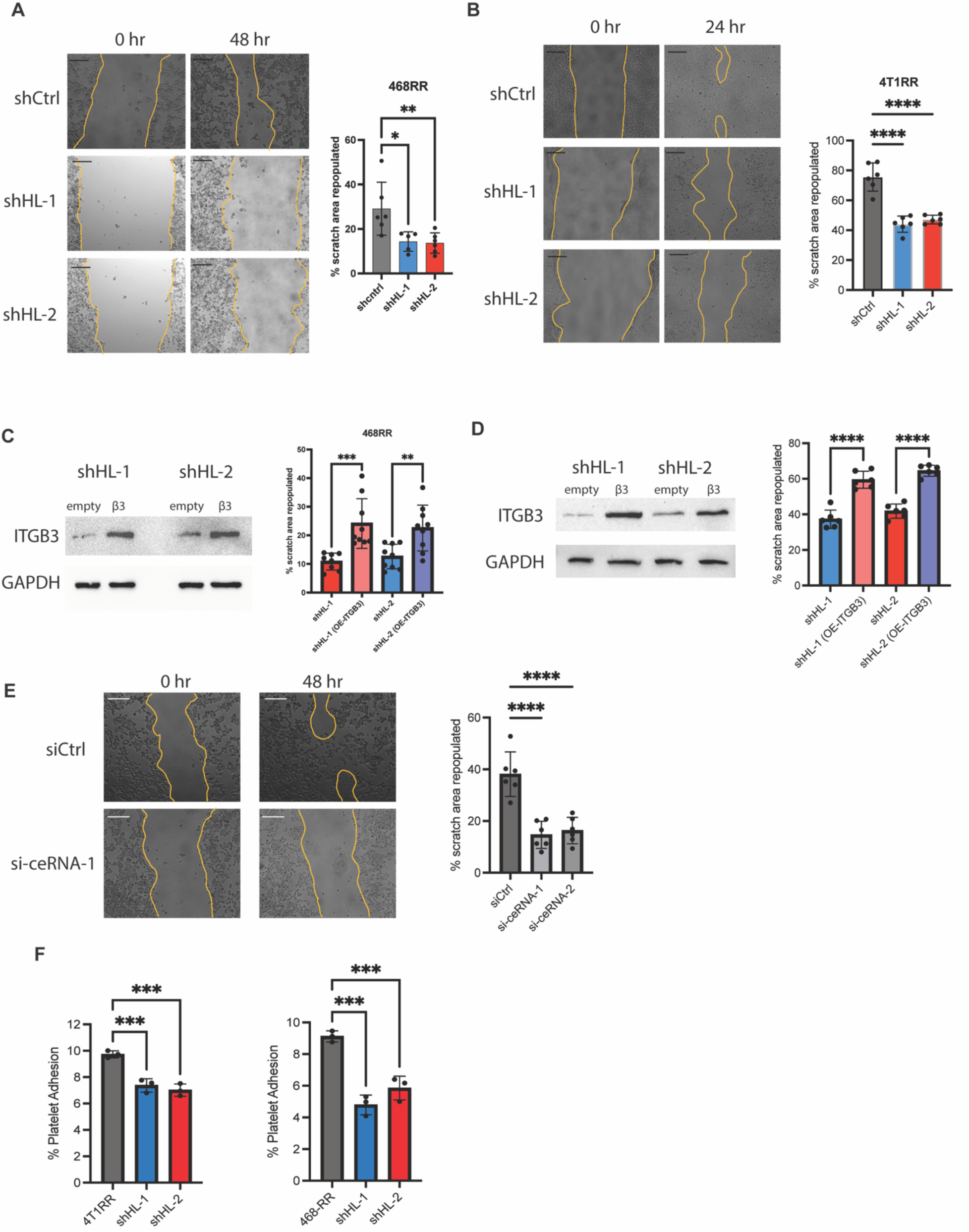
HNRNPL-derived circular RNAs drive migration by stabilizing ITGβ3 mRNA. **A.** The scratch area repopulated by MDA-MB-468-RR cells and the HNRNPL-knockdown cells after 48 hours. Scale bar is 200um. (n=6, mean ± SD, one-way ANOVA with Dunnett’s multiple comparisons tests). **B.** The scratch area repopulated by 4T1-RR cells and the HNRNPL-knockdown cells after 24 hours. Scale bar is 200um. (n=6, mean ± SD, one-way ANOVA with Dunnett’s multiple comparisons tests). **C.** The protein expression of MDA-MB-468-RR HNRNPL-knockdown cells given an empty vector or ITGβ3-GFP plasmid with the scratch area repopulated after 48 hours. (n=9, mean ± SD, unpaired t-test two-tailed) **D.** The protein expression of 4T1-RR HNRNPL-knockdown cells given an empty vector or ITGβ3-GFP plasmid with the scratch area repopulated after 24 hours. (n=6, mean ± SD, unpaired t-test two-tailed) **E.** The scratch area repopulated in 48 hours by MDA-MB-468RR cells given siCtrl, si-ceRNA-1, and si-ceRNA-2. The scale bar is 200um. (n=6, mean ± SD, one-way ANOVA with Dunnett’s multiple comparisons tests) **F.** The percent of platelets that were able to bind to the MDA-MB-468-RR and 4T1-RR cell lines along with their respective HNRNPL-knockdown cell lines seen *in vitro*. (n=3, mean ± SD, one-way ANOVA with Dunnett’s multiple comparisons tests).

## Discussion

The data obtained in this study using TNBC as a model advance the hypothesis that tumor cells that are resistant to radiation have a propensity to metastasize, an observation that strengthens the link between therapy resistance and the subsequent manifestation of aggressive behavior. We also demonstrate the mechanism for this phenomenon involves alterations in RNA metabolism that culminate in the enhanced expression of integrin β3, which drives metastasis of radioresistant cells. Specifically, the constitutive activation of Nrf2 promotes the expression of HNRNPL, which drives the expression of circular RNAs that could act as ceRNAs for let-7 miRNAs that mediate ITGβ3 degradation. The role of either circular RNAs or mRNA stability in radiation resistance has not been investigated extensively and our work provides significant advancement into their functional role of this pathway in enhancing the metastatic spread of therapy resistant TNBC.

A key finding in this study is our implication of RNA metabolism in radiation therapy resistance and, consequently, metastasis. Our work delineates two aspects of RNA metabolism in radioresistant cell lines: circular RNA formation and transcript stability. Circular RNAs not only serve as valuable biomarkers for cancer detection (39, 40) and progression (41, 42) because of their high stability and unique structure, but they also provide the functional capacity to drive diverse phenotypes in cancer by becoming ceRNAs or a hub for RNA binding proteins. Current work on HNRNPL-derived circular RNAs has demonstrated their relevance in several aspects of cancer ranging from tumor initiation (31) to response to immunotherapy (43, 44). We describe the contribution of three circular RNAs, which had not been elucidated prior to this study, in regulating integrin β3 migration in radioresistant TNBC. The concept of circular RNAs functioning as ceRNAs has been questioned especially in the context of one circular RNA having significant impacts on miRNA distribution and function (45–47). Importantly, however, our data support their ability to function as ceRNAs and regulate miR function. Specifically, we demonstrate that three circular RNAs (*circRAB12, circBACE2, circBIRC6*) have several binding spots for let-7 and they work in tandem to promote ITGβ3 mRNA stability within radioresistant TNBC. Our study focused on specific circRNAs that effect ITGβ3 transcript stability, but our use of IMCR-seq provides several other circular RNAs that may be important contributors to the aggressive nature of radioresistant TNBC. Current research in HNRNPL-derived circular RNAs is expanding and further work identifying these circular RNAs and their relative functions could uncover their contribution to cancer progression and its vulnerabilities.

Our findings provide insight into how integrin β3 is regulated in cancer, an issue that is significant and timely because pioneering studies by Cheresh (48, 49) and others (50) have highlighted the importance of this integrin in tumor stemness and resistance to chemotherapy, as well as its potential as a therapeutic target. A compounding factor, however, is that the ability of integrin β3 to contribute to these functions is independent of its ligand binding (48), which precludes the use of function-blocking Abs or other antagonists as possible therapeutics. Moreover, the available integrin β3 antagonists have not improved clinical outcomes for cancer patients and several adverse effects have been reported with their use (51, 52). As discussed by Cheresh (48, 49), a better understanding of how the ITGβ3 gene is regulated is needed to realize its therapeutic potential. Our work contributes to such an understanding by demonstrating that the stability of ITGβ3 mRNA mediated by an HNRNPL/circRNA mechanism is critical for the dynamic regulation of integrin β3, especially in response to the therapeutic stress of radiation. In the context of radiotherapy, previous studies have reported that increased surface expression of integrin β3 upon radiation of tumors provides the necessary downstream signaling to inhibit apoptosis (18) and promote DNA repair (53). Moreover, the current literature, as well as our data, delineate integrin β3 surface expression as a driver of platelet-tumor cell clusters that mediate extravasation and colonization within bone tissues (22, 54, 55).

The post-transcriptional regulation of the cancer transcriptome by RNA-binding proteins (RBPs) and miRNAs defines important aspects of tumor signaling. Previous work has demonstrated that the mRNAs that encode cancer initiation (56), epithelial-to-mesenchymal transition (57), stemness (58), and chemoresistance (59) are modulated by post-transcriptional regulators. Our data shows that the integrin β3-mediated migration and invasion of radioresistant TNBC is driven by HNRNPL, a key contributor to RNA stability mechanisms. Among the family of heterogenous nuclear ribonucleoproteins, HNRNPL is one of the few for which increased expression is associated with a worse overall survival in invasive breast carcinoma (60). Although one of its canonical functions is alternative splicing; we didn’t observe direct binding of HNRNPL to ITGβ3. Instead, the role of HNRNPL-mediated circular RNA formation is the mediator of ITGβ3 transcript stability in radioresistant cell lines. Our finding of an upstream regulator of integrin β3 expression that is dependent on ITGβ3 transcript stability provides an alternative mechanism by which cancer cells modulate this integrin. Moreover, uncovering the role of RNA metabolism in the context of radiation-resistant metastasis provides an opportunity to explore therapeutic vulnerabilities that exist in this pathway and could promote radiosensitivity.

The transcription factors that regulate HNRNPL transcription are not well established, and our work uncovers NRF2 activation as an upstream mediator of its expression. The incorporation of publicly available ChIP-seq data and our RNA sequencing data provided a few candidates, which we screened by downregulating their expression. The data shown in Fig 4 demonstrate NRF2 inhibition by pharmacological treatment or targeted RNA degradation decreased HNRNPL expression. We also demonstrated targeted NRF2 binding to the promoter region of HNRNPL by ChIP-qPCR. The current literature on NRF2, provided by us (61) and others (62–64), has provided extensive support for its role in mediating radiotherapy resistance by mediating ROS levels as well as inducing DNA repair pathways. However, its relevance in RNA metabolism pathways is still in its infancy. Further research on the RBPs that are mediated by NRF2 activation would be valuable in understanding their contribution to NRF2-driven cancer initiation and progression.

The cellular programs that are responsible for the enhanced metastatic potential of breast cancer often overlap with the ones that mediate therapeutic resistance. Our work establishes a factor of radiation resistance that mediates enhanced metastasis to the bone through mRNA stabilization. With advancements in technologies that capture circulating tumor cells, we propose that quantifying expression of HNRNPL and its circular RNA derivatives may be an effective biomarker of recurrence in TNBC patients given radiotherapy. Taking it one step further, therapeutic strategies that inhibit circular RNA formation via HNRNPL upon radiotherapy may decrease distant recurrence in breast cancer patients.

## Materials and Methods

### Sex as a biological variable

We implemented female mice in our experiments given that human breast cancer largely affects female patients.

### Cell Culture

The MDA-MB-468 human breast cancer cell line and the 4T1 mouse breast cancer cell line were purchased from American Type Culture Collection and were authenticated by the University of Arizona Genetic Core (UAGC). We developed radioresistant models of MDA-MB-468 and 4T1 by giving a total of 50 Gy over the course of 6-8 weeks using the following treatment schedule: 2Gyx5, 4Gyx3, 6Gyx3, and 10Gyx1 (17). Before moving on to the next radiation dose, we waited for the cells to reach 70-80% confluency in the plate.

### Reagents

Propidium iodide was purchased from Thermo Fisher Scientific (P1304MP). Annexin V-FITC was purchased from Invitrogen (A13199). Mitomycin C was purchased from Millipore Sigma (10107409001). Matrigel Growth Factor Reduced (GFR) Basement Membrane Matrix was purchased from Corning (354230). The NRF2 inhibitor, ML385, was ordered from MedChemExpress (HY-100523); the NRF2 inducer, Dimethyl Fumarate was ordered from Sigma (242926). The following antibodies were used for immunoblotting: GAPDH (14C10) [Cell Signaling Technology 2118S], beta Actin (8H10D10) [Cell Signaling Technology 3700S], alpha Tubulin (TU-02) [Santa Cruz Biotechnology sc-8035], anti-FLAG (M2) [Sigma Aldrich F3165] HNRNPL (4D11) [Novus Biologicals NB120-6106SS], ITGβ3 [Cell Signaling Technology 4702S]. The following Abs were used for flow cytometry: ITGβ3 (D-11) [Santa Cruz Biotechnology sc-36567], anti-Mouse-Alexa Fluor 647 [Invitrogen A-21235]. For immunofluorescence the following antibodies were used: NRF2 (D1Z9C) [Cell Signaling Technology 12721s] and anti-rabbit-AF488 [Invitrogen A-11008].

### Constructs

The following lentiviral shRNAs were obtained from our core facility: HNRNPL (TRCN0000017243-human, TRCN0000017245-human, TRCN0000112038 - mouse, TRCN0000112039 - mouse), NRF2 (TRCN0000007556 and TRCN0000007557). shCtrl vectors were pLKO scramble shRNAs (Addgene, 1864). The following siRNAs were designed and ordered from IDT: mouse siITGβ3-1, mouse siITGβ3-2, sicircBIRC6-1, sicircBIRC6-1, sicircBACE2-1, sicircBACE2-2, sicircRAB12-1, sicircRAB12-2. The sequence of the siRNAs is provided in Supplementary Table 2. Control siRNA-A (sc-37007, Santa Cruz). Lentivirus packaging vectors were obtained from Addgene pMD2.G (12259) and psPAX2 (12260). A lentiviral plasmid expressing ITGβ3-GFP was obtained from Addgene (205091). A HNRNPL-FLAG tagged construct was ordered from Sino Biological (HG18369-CF). A lentiviral plasmid that can form circular RNA (pF CAG mc2 IRES neo) was obtained from Addgene (206232). A gBlock of *circRAB12* was designed and ordered from IDT. The circular RNA forming plasmid was digested with BsmbI-v2 (R0739S, NEB) and then cloned with the *circRAB12* gBlock using the 2x HiFi DNA Assembly Master mix (E2621S) based on the manufacturer’s instructions. The empty plasmid control for these experiments was the pF CAG mc2 IRES neo plasmid.

### Clonogenic assays

Cells were treated with a radiation dose ranging from 0-6Gy and then a predetermined number of cells was added to each well and evenly distributed based on cell type and radiation dose. The media in the wells was replaced every 3 days. After 10-14 days, the plates were fixed with 4% paraformaldehyde for 15 minutes, washed three times with 1X PBS, and stained with 0.5% crystal violet in 80% methanol for 45 minutes. Colonies with more than 50 cells were counted. The plating efficiency was calculated as the number of colonies formed divided by the number of cells added to the well for the non-irradiated control. The surviving fraction was calculated based on the ratio of colonies in the treatment group to the number of colonies in the non-irradiated samples and the plating efficiency of the cell line. The radiosensitivity enhancement ratio was calculated by measuring the area under the curve from the clonogenic assay of the control to the experimental group.

### Flow cytometry

To assess surface expression of ITGβ3 in cell lines, we incubated 1×10^6^ cells in 100μL of PBS with primary antibody at a concentration of 1:50 for 30 min on ice. The cells were washed with PBS and centrifuged for 3 min at 300g, which was repeated for a total of three times. After the last wash, the cells were resuspended in 100μL of PBS and incubated with secondary antibody at a concentration of 1:1000. Apoptosis after irradiation was assessed using Annexin V binding to the cell surface. The cells were collected and resuspended in annexin-binding buffer (10 mM HEPES, 140 mM NaCl, and 2.5 mM CaCl_2_, pH 7.4) and aliquoted in a glass tube to have 1×10^5^ cells in 100μL. 5μL of Annexin V-FITC was added to the solution along with 3μl of 100μM solution of PI. The samples were incubated in the dark for 15 minutes. An additional 400μL of the annexin-binding buffer was added to the solution and then placed on ice until analyzed on the flow cytometer (BD FACS Celesta).

### Immunoblotting

Protein extraction was done by scraping cells on ice with RIPA buffer (BP-115DG, Boston Bioproducts) supplemented with protease and phosphatase inhibitors (Thermo, 78442). Subsequently, laemmli buffer (BP-111R, Boston Bioproducts) was added to each sample and the lysate was boiled and separated using SDS-PAGE. Immunoblotting primary antibodies were used at the following concentrations: HNRNPL 1:2000, integrin β3 1:1000, Tubulin and GAPDH 1:2000, and anti-FLAG 1:1000. The secondary antibodies conjugated with HRP were used at a concentration of 1:5000.

### mRNA quantification

mRNA quantification was completed by first extracting RNA using the NucleoSpin RNA kit (Macherey-Nagel 740955.50) and proceeding to cDNA synthesis (AZ-1996, Azura Genomics). The relative expression levels were quantified using the Azura View GreenFast qPCR Blue Mix LR master mix (AZ-2320, Azura Genomics). Experiments were performed in triplicate and normalized to *GAPDH*. The primers used are in Supplementary Table 3

### RNA-seq

RNA was extracted from the indicated cells using a NucleoSpin RNA kit (740955.50 Macherey-Nagel) and sent to Novogene (4T1-Par vs 4T1-RR and MDA-MB-468 vs MDA-MB-468-RR) for sequencing. Library for RNA-Seq was prepared according to KAPA Stranded mRNA Hyper prep polyA selected kit with 201–300 bp insert size (KAPA Biosystems, Wilmington, MA) using 250 ng total RNAs as input. Final library quality and quantity was analyzed by Agilent Technologies 4200 station and Qubit 3.0 (Thermo Fisher Scientific Inc, Waltham, MA) Fluorometer. 150 bp paired-end reads were sequenced on Illumina HiseqX (Illumnia Inc., San Diego, CA). Each sample had a sequencing depth of 30–40 million. RNASeq analysis was performed with OneStopRNAseq workflow (65). Paired-end reads were aligned to human primary genome T2T-CHM13v2.0, with star_2.5.3a (66). Aligned exon fragments with mapping quality higher than 20 were counted toward gene expression with featureCounts_1.5.2 (67). Differential expression (DE) analysis was performed with DESeq2. Significant DE genes (DEGs) were filtered with the criteria FDR <0.05.

### Induro-Reverse Transcriptase mediated circular RNA sequencing (IMCR-seq)

Total RNA from the cell lines (MDA-MB-468-RR shctrl and shHNRNPL-1) were extracted using a NucleoSpin RNA kit (740955.50 Macherey-Nagel). To sequence the circular RNA, the following protocol was adapted from the IMCR-seq protocol recently published (). A total of 4 tubes were taken and each tube contained: 500ng of total RNA (max 6uL), 2 uL 100 uM random N6 (NEB #S1230S), and 2 uL ddH2O. The tubes were incubated at 72°C for 5 minutes then cooled down to 4°C with 0.1°C/s ramp. Each tube was then taken and the following were added to reach a total volume of 40 uL: 8 ul 5X Induro Buffer, 4.8 ul MgCl2 (NEB B9021S: 25 mM), 2 ul 10 mM dNTPs, 2 ul Induro RT (NEB M0681: 200 units/ul), and 13.2 ul ddH2O. The tubes were incubated at: 23°C for 10 minutes, 30°C for 5 minutes, warmed up to 55°C with 0.1°C/sec ramp, 55°C for 1 hour, and 95°C for 1 minute to heat inactivate the RT. All four tubes were collected for a total volume of 160 uL and then mixed with 40 uL water and 200 uL AMPure beads (A63880, Beckman Coulter). The tubes were put on a rotator for 10 minutes. The tubes were placed on a magnetic rack to remove the liquid on the magnet and then resuspended in 300 uL 10mM Tris-HCl pH:8.0, with 72 uL Mg500PEG5 solution (500mM MgCl and 5% PEG8000 in 10mM Tris-HCl pH:8.0). The tubes were on the rotator for 30 minutes, placed on magnetic rack to remove the liquid, washed with fresh 80% EtOH and eluted the cDNA in 33 uL water. A 5uL aliquot of the cDNA was taken and mixed with 5 ul 10X Phi29 Buffer, 5 ul 10mM dNTPs, 2.5 ul 1mM Exo-resistant N6 primers (SO181 Thermo Scientific), 2 ul phi29 (M0269 NEB), and 30.5 ul ddH2O. This was done for a total of 6 tubes and all of them were incubated at 30°C for 6 hours. All tubes were combined together along with 15 ul NEBuffer 2 (B7002S NEB), 100 uL ddH2O, and 75 uL T7 Endonuclease I (M0302S NEB). The tube is incubated at 37°C for 1 hour. 500 uL of AMPure beads will be added to the mixture and incubated on the rotator for 10 minutes. The liquid is removed by placing the tube on the magnetic stand and the beads are mixed with 300 ul 10mM Tris-HCl pH:8.0 and 72 ul Mg500PEG5 solution (500mM MgCl and 5% PEG8000 in 10mM Tris-HCl pH:8.0). The tube is placed on the rotator for 30 minutes. Lastly the liquid is removed with the magnetic stand, beads are washed with fresh 80% EtOH twice, and finally eluted in 60ul water.

### Library Preparation, Nanopore Sequencing, and circular RNA analysis

1.5ug of DNA from IMCR-seq was used for library preparation using Nanopore ligation sequencing kit (SQK-LSK114) according to the manufacturer’s protocol, with the following specific alterations.. During library preparation, pipetting of DNA samples was avoided to minimize shearing of long DNA. Incubation time for all steps was extended by 5-10 minutes. For last Ampure clean-up set, 100uL was used. Long Fragment Buffer was used twice to wash. And all elutions were done at 55C. All libraries were sequenced on R10 flow cell on MinION for 24 hours with a minimum read cutoff of 1 kilobase. High-accuracy basecalling was done on MinION or using dorado version 0.8.0. Passed reads or all reads were used as an input for CIRI-long (68) using human genome assembly GRCh38.p14 and GENCODE comprehensive gene annotations to identify circRNAs. CircRNA isoforms identified from all replicates were collapsed and counted using CIRI-long collapse program. circRNA isoforms with an average CPM of less than 3 in either condition were filtered out and summed at the gene-level. Gene-level circRNA counts were used for DESeq2 (69) with a default parameter to identify differentially expressed circRNAs. Top 5 genes based on log2 fold-change * -log10(FDR) in either direction were plotted.

### Analyzing circular RNAs that are ceRNAs

We first filtered the circular RNAs that were greater than 2-fold increase in the MDA-MB-468-RR cells compared to the HNRNPL knockdown cells and had a greater than 3 CPM value. We focused our analysis on let-7 microRNAs and used those sequences along with the circular RNA sequences as input for Circr (33), a computational program that predicts circular RNA and miRNA binding. To predict the capacity of the circular RNAs that function as an efficient sponge for let-7, we developed a score that considers the number of binding sites for let-7 miRNAs and the differential expression compared to the HNRPNL knockdown cells. With this score, we were able to identify the top circRNAs that could function as ceRNAs for enhanced ITGβ3 mRNA stability.

### Invasion Assay

For invasion analysis, 30ug of Matrigel was added to a 24-well transwell membrane and allowed to harden at 37°C for 2 hours. The upper part of the well had 3×10^4^ cells in serum-free media and the lower part of the chamber had 500ul of conditioned media. The transwells were placed in a DAPI solution for 10 minutes, fixed to a coverslip, and imaged using a microscope. A total of 5 images were taken from each transwell to count the total number of cells that migrated. This experiment was completed in triplicate.

### Scratch-wound migration assay

Cells were plated on a 24-well plate at a density of 75,000 cells/well for the 4T1 cell lines and 2×105cells/well for the MDA-MB-468 cell lines and allowed to attach overnight and reach a confluency of ∼80-90%. After reaching the confluency criteria, the cells were pretreated with mitomycin c (10107409001 Roche) at 5uM concentration for 1 hour followed by three washes of 1x PBS. A scratch was made in the well using a 20ul pipette tip and the well was washed with 1x PBS prior to imaging. The MDA-MB-468 cell lines were cultured in DMEM/F12 media + 2% FBS whereas the 4T1 cell lines were cultured in RPMI media + 0.5% FBS. The wells were taken in the microscope to image the scratches at 48 hours for the MDA-MB-468 cell lines and 24 hours for the 4T1 cell lines.

### Platelet adhesion assay *in vitro*

The following protocol was adapted from Egan et. al to look at platelet adhesion to our cell lines (70). The cell lines were plated on a 96-well plate at a concentration of 1×10^4^ cells/well for 4T1 cell lines and 5×10^4^ cells/well for MDA-MB-468 cell lines and incubated in 37°C incubator overnight. The platelets were ordered from Human Cells Bio (Platelet-F5B, Donor HH060), and centrifuged with Acid-Citrate-Dextrose [ACD: 38 mM citric acid, 75 mM sodium citrate, 124 mM D-glucose] and PGE_1_ [1 μM] at 720g for 10 minutes. The supernatant was removed, and platelets were resuspended in JNL buffer [130 mM NaCl, 10 mM sodium citrate, 9 mM NaHCO_3_, 6 mM D-glucose, and 0.9 mM MgCl_2_, 0.81 mM KH_2_PO_4_, and 10 mM Tris, pH 7.4] supplemented with 1.8mM CaCl_2_ at a concentration of 1.5×10^8^ cells/mL. Platelets in suspension were incubated with Calcein AM (Cayman Chemical, 14948) at a concentration of 2μg/mL and then 100μL of the suspension was added in each well. The total fluorescence as measured using a GloMax plate reader [Promega, 485/535 nm]. The plate was washed three times with 100μL JNL buffer. After the final wash, 100μL of JNL buffer was added to the well and fluorescence was quantified using the plate reader. The % platelet adhesion was quantified as [remaining fluorescence – blank]/[total fluorescence – blank] ×100.

### RNA stability

Cells were treated with 5ug/mL of actinomycin D for 0, 1 2, or 4 hours and then total RNA was extracted using the NucleoSpin RNA kit. Reverse transcription of the samples was carried out using the Azura cDNA synthesis kit (AZ-1995). Quantitative PCR was done using the ITGβ3 primers and SYBR Green Master mix.

### Immunofluorescence microscopy

For cell imaging, cells were cultured on 35mm glass bottom dishes, washed with PBS, fixed with 4% paraformaldehyde in PBS for 15 min at room temperature. Subsequently, cells were washed with PBS, put in blocking buffer (5% normal goat serum (Sigma-Aldrich, G9023), 0.3% Triton X-100 in PBS) for 1 hour, and then incubated with primary antibodies in antibody dilution buffer (1% BSA, 0.3% Triton X-100 in PBS) overnight at 4^ο^C. Cells were washed with PBS three times followed by 1 hour incubation with secondary antibodies and DAPI (1:1000, Invitrogen). The cells were mounted in a 0.1M *n*-propyl gallate, 90% (by volume) glycerol, and 10% PBS solution. To assess the nuclear localization of NRF2, we developed masks for the nuclear region and whole cell and quantified the total fluorescent intensity in the field.

### ChiP

We used the ChIP-IT Express Chromatin Immunoprecipitation kit (Active Motif). The following primers (5’-GGACTGTTGATACGCTCTGATT-3’, 5’-GTATTTGCGCATGCGTTCTC-3’) amplified the region of the *HNRNPL* promoter with a NRF2 peak.

### RNA Immunoprecipitation

The following protocol was an adapted version of a previously published in vitro RNA immunoprecipitation (iv-RIP) protocol (71). The protein from the cells were collected from a 10cm plate by adding 300ul of RIPA buffer and using a rubber scrapper. The lysate was transferred to an Eppendorf tube and put on rotator for 30 min at 4°C. The sample was then sonicated using a Bioruptor with 4 cycles of 5s ON and 30s OFF. The lysate was centrifuged at 12000g for 15 min at 4°C. The supernatant was collected and transferred to a new tube. The protein concentration of the lysate was taken and 1mg of protein was mixed with 1ug of HNRNPL antibody or IgG antibody. The tube was placed on a rotator overnight at 4°C. The protein-antibody complex was incubated with 10uL of protein G magnetic beads that were pre-washed with lysis buffer. The mixture was kept on a rotator for 4 hours at 4°C. The tube was kept on a magnet, supernatant removed, washed with lysis buffer twice, and then resuspended in RIPA buffer. The resuspended beads-antibody-protein complex is mixed with 1ug of total RNA on a rotator at room temperature for 45 minutes. The tube was kept on a magnet, supernatant removed, washed with RIPA buffer twice, and then resuspended in 50ul RNAse-free water.

### Mouse experiments

To evaluate spontaneous metastasis between 4T1 and 4T1-RR, the cells were labeled with luciferase using a lentivirus and antibiotic selection. A total of 1×10^5^ cells were orthotopically injected into the mammary fat pad of BALB/C mice purchased from Charles River Laboratories. The tumors were measured using calipers and the volume was calculated using the following equation: height x width^2^/2. Bioluminescent images of the thoracic region of the mouse were taken using the Perkin Elmer IVIS Spectrum CT and evaluated for metastasis by quantifying the flux (photon/s). To study hematogenous metastasis capacity of MDA-MB-468 and MDA-MB-468-RR, the cells were labeled with luciferase using a lentivirus followed by antibiotic selection. An intracardiac injection of 2×10^5^ cells were delivered to the circulatory system of NSG mice (NOD.Cg-Prkdc*^scid^*Il2rg*^tm1Wjl^*/SzJNSG purchased from Jackson Laboratory) and bioluminescent images were taken to determine organotropic differences in metastasis.

### Clinical Patient Data

The patient data used in this study were taken from cBioPortal (72–74) for Cancer Genomics, which has access to breast cancer patient data retrieved from The Cancer Genome Atlas Program (TCGA), or using TIMER2.0 (75), which allows for correlation of gene expression for basal-subtype breast cancer patients.

### Data Availability

The RNA-seq data have been deposited in the NBCI GEO database as GSE272692 (4T1 vs 4T1-RR), GSE289555 (MDA-MB-468 vs MDA-MB-468-RR), and GSE290005 (circular RNA in MDA-MB-468-RR vs HNRNPL knockdown). Values for all data points in graphs are reported in the Supporting data values file. Raw immunoblot data are reported in the full unedited blot and gel images file.

### Statistical Analysis

Student’s t test was used to compare between two groups and more than 2 groups was compared using either one-way ANOVA followed by Dunnett’s multiple comparisons test or 2-way ANOVA followed by Tukey’s multiple comparisons test. Kaplan-Meir analysis was completed using the Gehan-Breslow-Wilcoxon. All statistical tests were carried out using GraphPad Prism 10.0 with a significance level set at *P* less than 0.05 and the following symbols represent the associated P-value: \**P*<0.05, \*\**P* <0.01, *** *P*<0.001, and **** *P*<0.0001.

## Author Contributions

AK, HLG and AMM contributed to overall project direction, study design and data analysis. RL and LJZ performed the RNA-seq analysis. KK, BP and WAF provided advice, design, data collection and analysis for the IMCR-sequencing that identified circular RNAs. AK, CW and HLG performed the animal experiments. AK and AMM wrote the manuscript.

## Acknowledgements

We thank Dr. Matthew Hemming (Dept. of Medicine, UMASS Chan Medical School) and Dr. Athma Pai (RNA Therapeutics Institute, UMASS Chan Medical School) for their critical comments and suggestions on the manuscript. We would like to thank Dr. Mattia Forcato (Università Degli Studi Di Padova, Italy) for developing Circr and providing insight into using the computational tool. Lastly, Dr. Timothy FitzGibbons and Mark Kelly provided their expertise for the intracardiac injections of the cells into the mice. “The results shown in Figure 4B here are in part based upon data generated by the TCGA Research Network: https://www.cancer.gov/tcga.” This work was supported by NIH Grants R01 CA285607 (AMM), R50 CA221780 (HLG), and F30 CA275327-01A1 (AK).

## Supplementary Figures

**Supplementary Figure 1:**
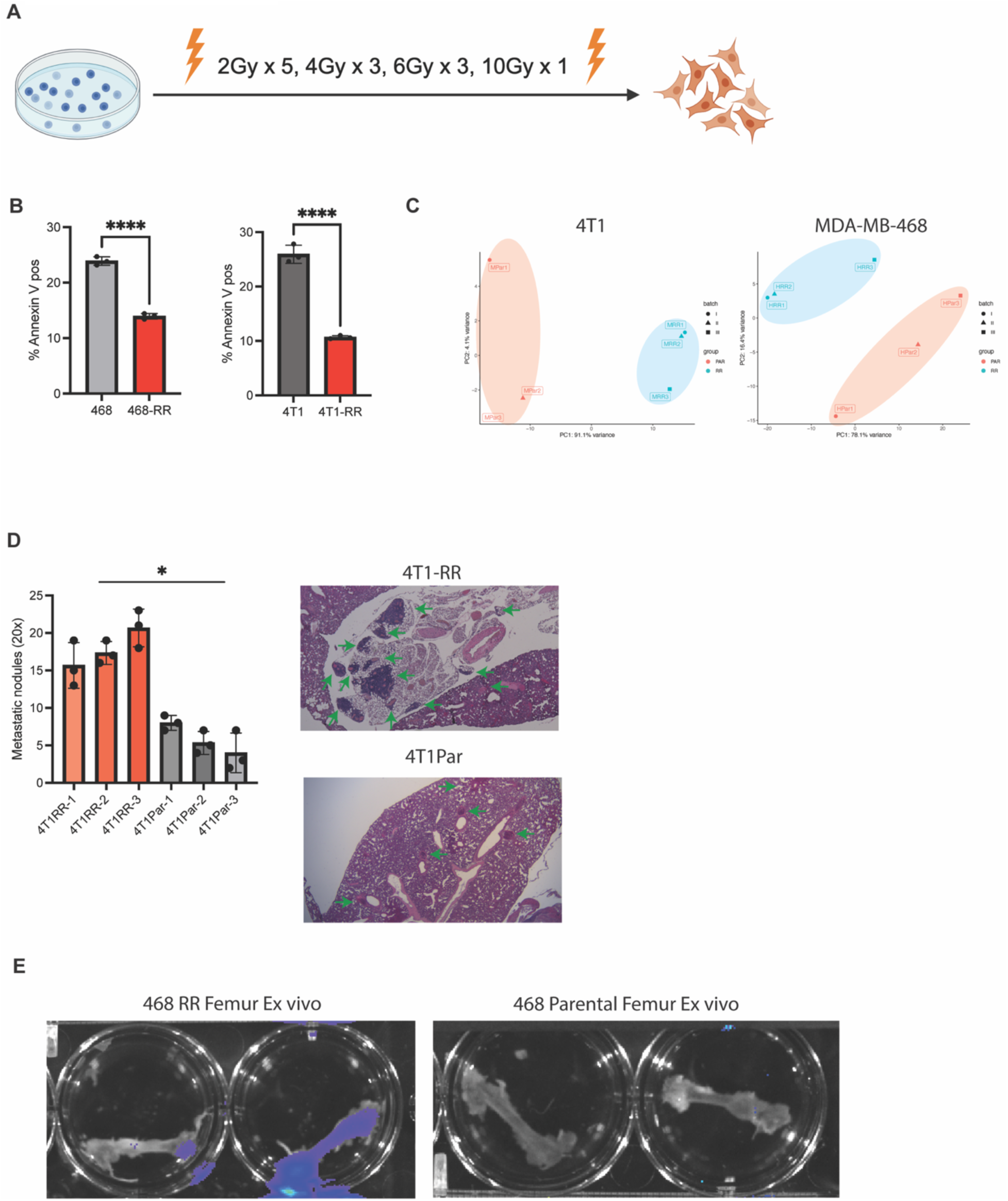
Characterizing radioresistant and parental cell lines. **A.** The diagram demonstrates the radiation dose escalation strategy that was implemented to develop the radioresistant cell lines. **B.** The percent of Annexin-V positive cells 48 hours after irradiation at a dose of 8Gy of the parental and radioresistant cell lines. n=3 independent samples. **C.** The PCA plots for the 4T1-RR, 4T1-parental, MDA-MB-468-RR, and MDA-MB0468-parental cell lines after differential gene expression analysis. **D.** The number of macro-metastatic sites observed in the thoracic cavity of the mice injected with either 4T1-RR or 4T1 cells. n=3 per section. **E.** The bioluminescent images of the femur ex vivo from the mice injected with MDA-MB-468-RR vs MDA-MB-468.

**Supplementary Figure 2:**
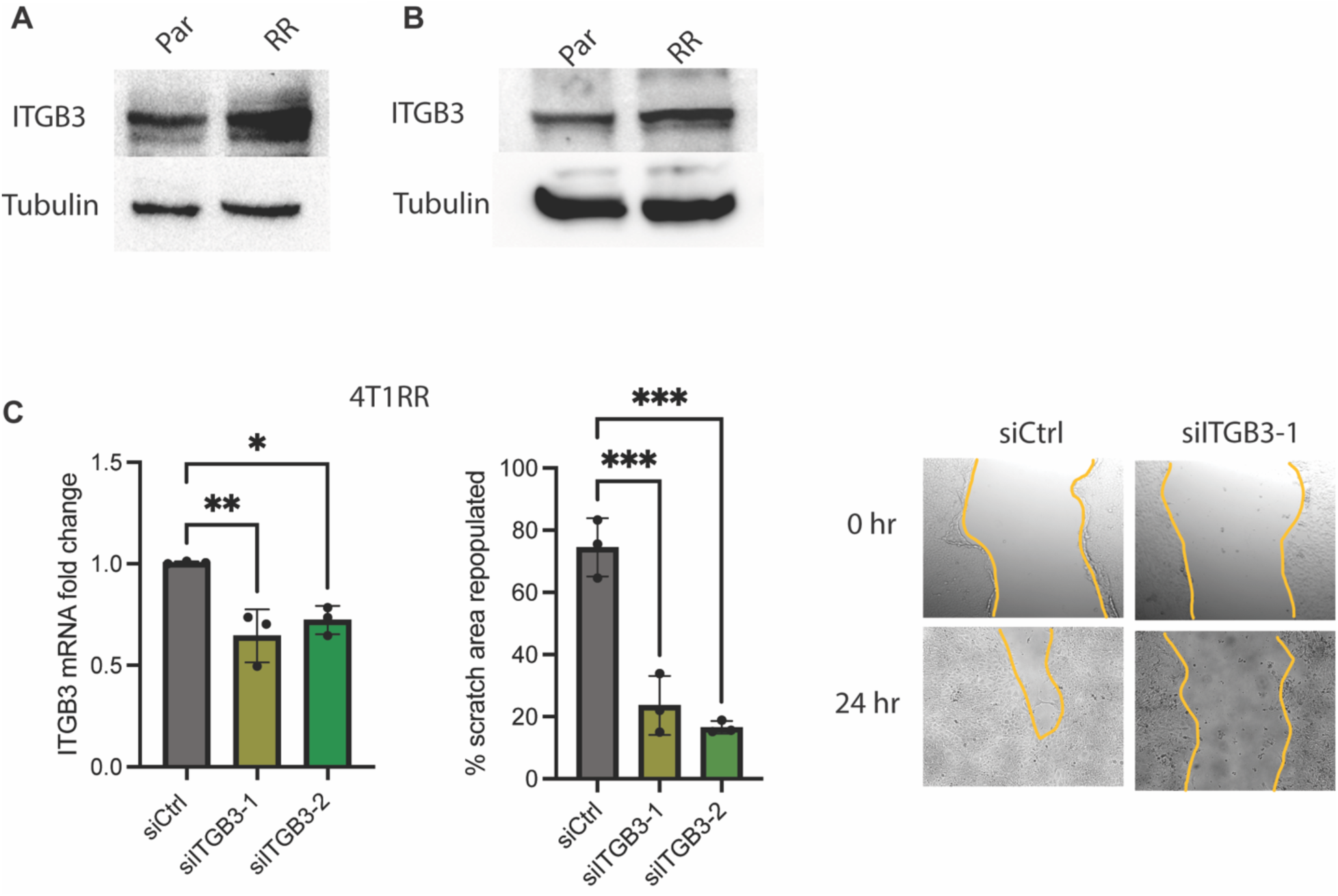
ITGβ3 expression and function in radioresistant TNBC. **A.** An immunoblot of ITGβ3 and tubulin from the MDA-MB-468-RR cells and the MDA-MB-468 cells. **B.** An immunoblot of ITGβ3 and GAPDH from the 4T1-RR cells and the 4T1 cells. **C.** The expression of ITGβ3 after treatment with siRNAs (siCtrl, siITGB3-1, and siITGB3-2) and the migration capacity as determined by the scratch-wound assay (The scale bar is 200um). Data are represented as the mean ± SD. The P values in panel **C** were obtained by on-way ANOVA followed by Dunnett’s multiple comparisons test.

**Supplementary Figure 3:**
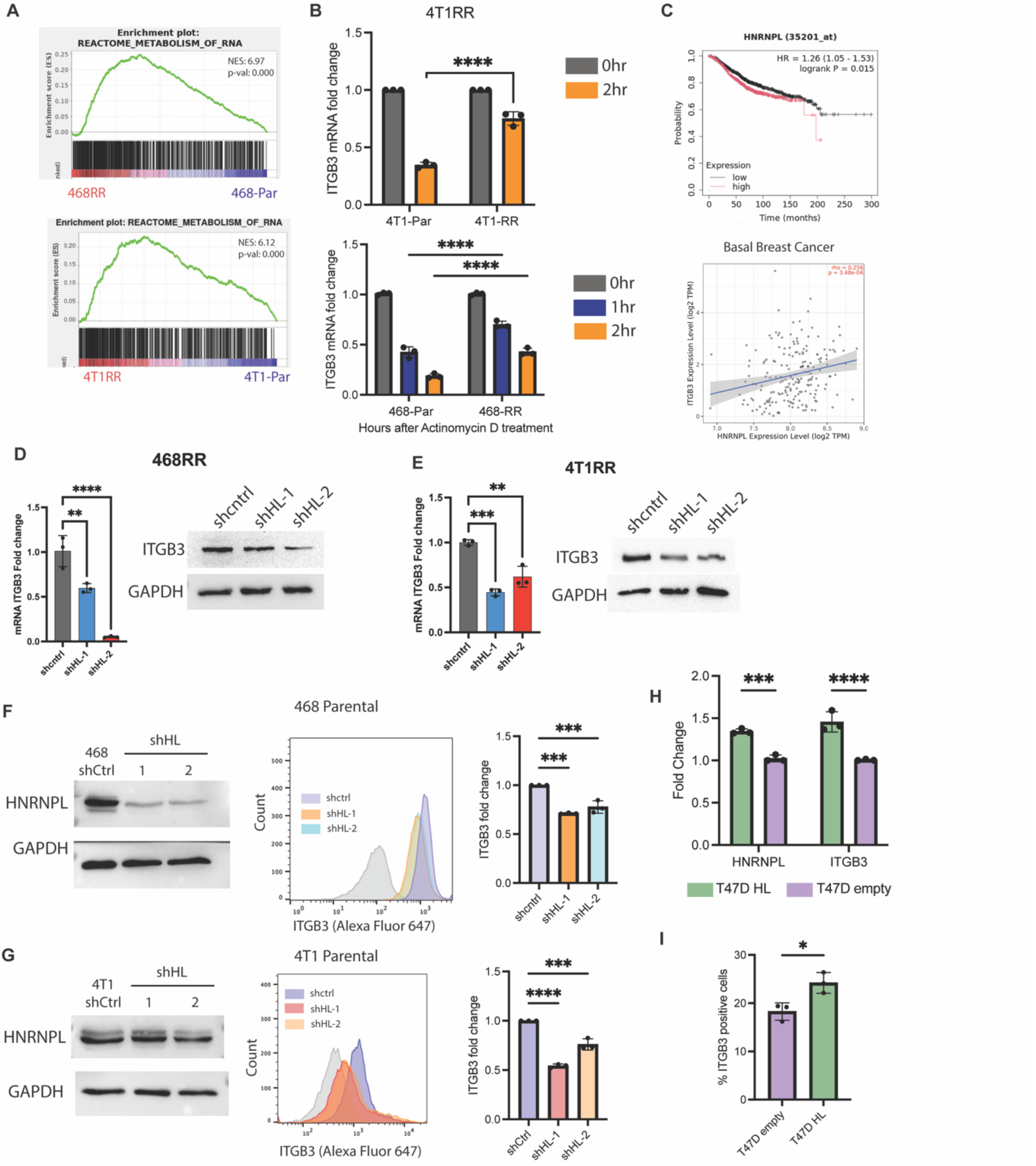
ITGβ3 mRNA stability is reliant on HNRNPL expression. **A.** The enrichment plot for the geneset “REACTOME METABOLSIM OF RNA” that is upregulated in the MDA-MB-468-RR cells and 4T1-RR cells. **B.** The transcript stability of ITGβ3 as determined by the expression of ITGβ3 after different time points (0, 1, or 2 hours) of actinomycin d treatment in the radioresistant cells and parental cells based on RT-qPCR and normalized to expression at the 0 hour timepoint. **C.** Kaplan-Meier overall survival for breast cancer patients segregated by median HNRNPL expression in tumors taken from KM plotter. The correlation between ITGβ3 and HNRNPL expression in Basal Breast Cancer. **D.** The mRNA and protein expression of ITGβ3 when diminishing HNRNPL expression in the MDA-MB-468-RR cells **E.** The mRNA and protein expression of integrin β3 when diminishing HNRNPL expression in the 4T1-RR cells. **F.** An immunblot of HNRNPL expression in the MDA-MB-468 cells and the HNRNPL knockdown cells along with the flow cytometry data showing ITGB3 surface expression. **G.** An immunblot of HNRNPL expression in the 4T1T cells and the HNRNPL knockdown cells along with the flow cytometry data showing integrin β3 surface expression. **H.** The mRNA expression of HNRNPL and ITGβ3 in the T47D cells transfected with empty vector of HNRNPL FLAG-tagged vector. **I.** The percent of cells expressing integrin β3 in the T47D cells transfected with empty vector of HNRNPL FLAG-tagged vector. Data are represented as mean ± SD. The P values in panel **B** were obtained from two-way ANOVA with Tukey’s multiple comparisons tests, panel **D**-**G** were obtained from one-way ANOVA with Dunnett’s multiple comparisons tests, panel **H** and **I** were obtained from student’s t-test two-tailed.

**Supplementary Figure 4:**
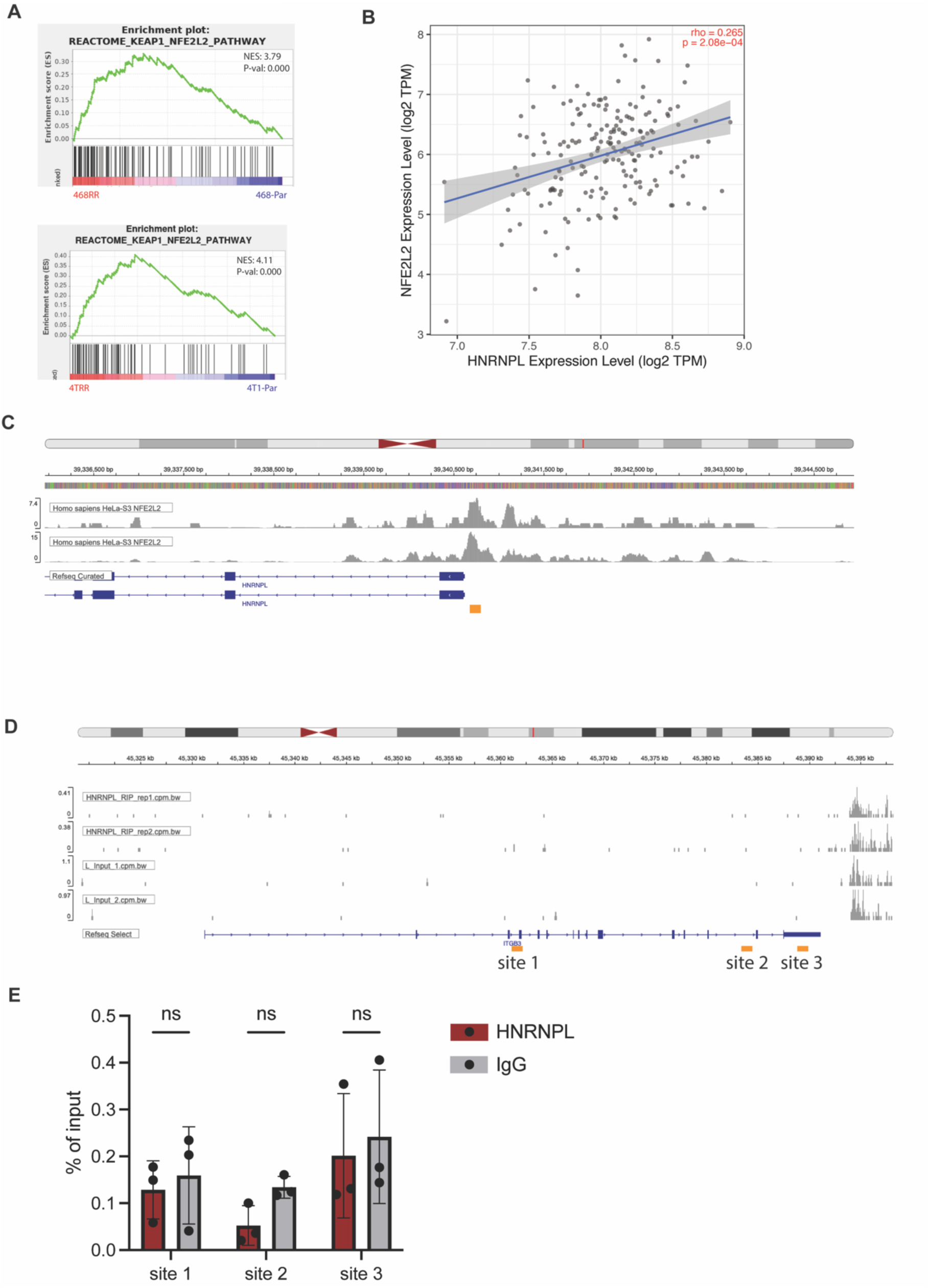
NRF2 binding at HNRNPL and lack of binding of HNRNPL to ITGβ3 transcript. **A.** The enrichment plot for the geneset “REACTOME KEAP1 NRF2 PATHWAY” that is upregulated in the MDA-MB-468-RR cells and 4T1-RR cells. **B.** The correlation between NRF2 expression and HNRNPL expression in basal breast cancer patients taken from TIMER2.0. **C.** The ChIP-seq data from HeLa cells identifying the binding sites of NRF2 near the HNRNPL locus from GSE91997. The orange box designates the region where primers were designed to detect NRF2 binding near the HNRNPL promoter. **D.** The RIP-seq data from prostate cancer cells identifying the potential binding sites of HNRNPL near the ITGβ3 transcript from GSE72844. **E.** The RIP-qPCR data of potential HNRNPL binding sites near ITGβ3 taken from the MDA-MB-468-RR cells. Data are represented as mean ± SD. The P values in panel **E** were obtained unpaired t-test two-tailed.

**Supplementary Figure 5:**
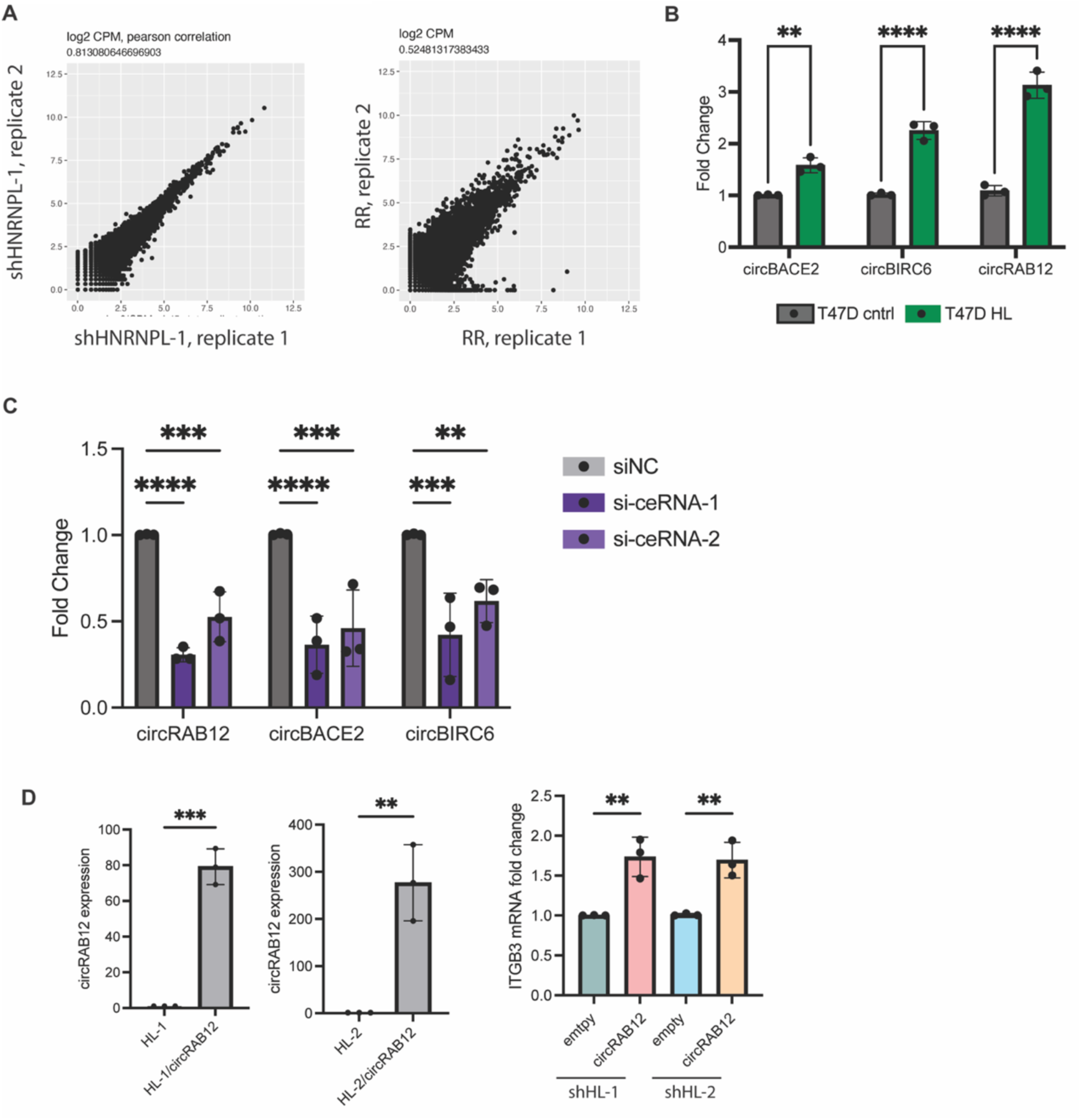
The role of circRNAs in mediating ITGβ3 expression. **A.** The Pearson correlation of the circular RNA isoforms for each biological replicate of the HNRNPL knockdown cells and RR cells. **B.** The expression of the ceRNAs based on RT-qPCR in T47D cells given empty vector or HNRNPL-FLAG plasmid. **C.** The expression of *circRAB12*, *circBACE2*, and *circBIRC6* in the control cells (siNC) compared to the siRNA knockdown cells (si-ceRNA-1 and si-ceRNA-2). **D.** The mRNA expression of *circRAB12* after transfection of the *circRAB12* or empty plasmid into the HNRNPL knockdown cells. The mRNA expression of ITGβ3 in those cells. Data are represented as mean ± SD. The P values in panel **B** and **D** were obtained unpaired t-test two-tailed, in panel **C** were obtained by one-way ANOVA with Dunnett’s multiple comparisons test.

---

**Supplementary Table 1:** The scoring system for the capacity of circRNAs to sponge let-7 miRNAS.

| circ Name | number of sites | RR_exp | HL_exp | gene | score |
| --- | --- | --- | --- | --- | --- |
| chr18:8624938-8638416 | 6 | 103.59782<br>1 | 26.205756<br>3 | RAB12 | 464.35239<br>1 |
| chr10:126970702-127061776 | 5 | 54.113264<br>1 | 25.706576<br>2 | DOCK1 | 142.03343<br>9 |
| chr1:108885275-108904254 | 5 | 44.109349<br>2 | 19.425905<br>2 | GPSM2 | 123.41722 |
| chr2:32377588-32395593 | 5 | 36.456271<br>3 | 12.883679<br>2 | BIRC6 | 117.86296<br>1 |
| chr22:20933779-20934244 | 5 | 38.959292<br>6 | 16.193917<br>8 | CRKL | 113.82687<br>4 |
| chr16:81351583-81377702 | 4 | 43.645567<br>4 | 17.975267<br>9 | GAN | 102.68119<br>8 |
| chr7:24623666-24680520 | 5 | 34.271624<br>7 | 14.480768<br>2 | PALS2 | 98.954282<br>6 |
| chr21:41226266-41257326 | 4 | 32.454411<br>7 | 10.261428<br>5 | BACE2 | 88.771933 |

**Supplementary Table 2:**
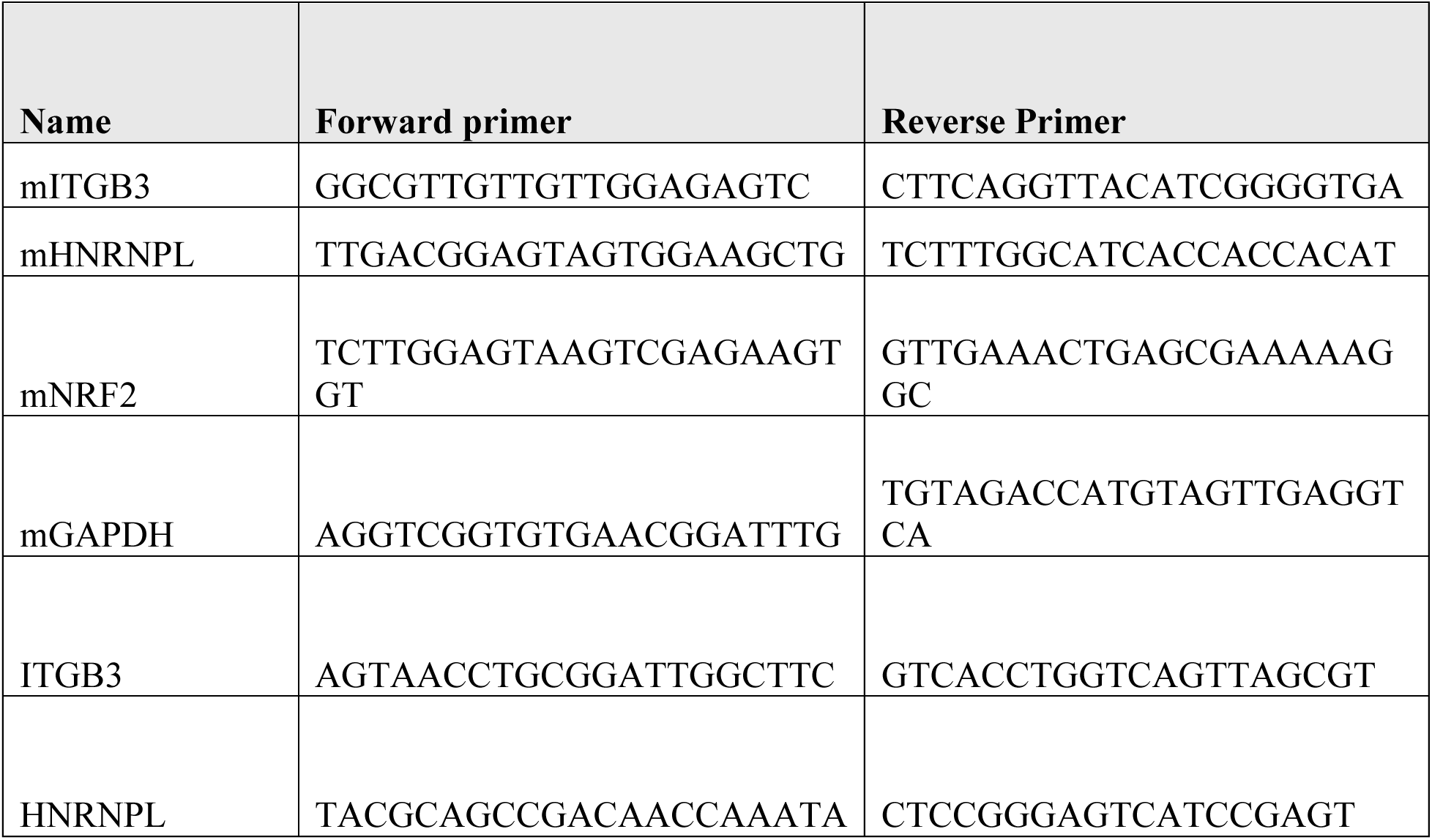
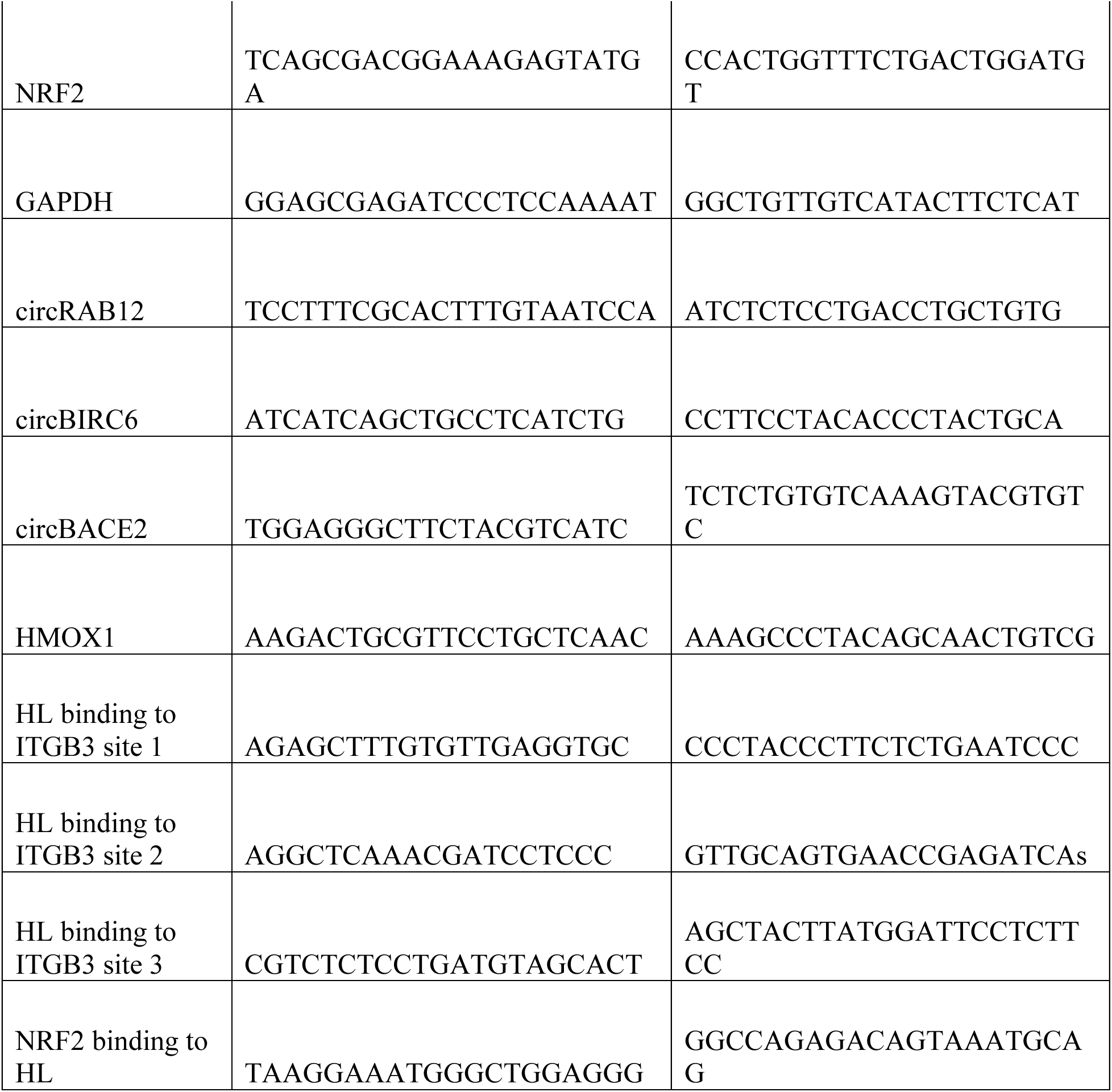
The primers used for RT-qPCR, ChIP-qPCR, and RIP-qPCR.

**Supplementary Table 3:** The siRNA sequences.

| siRNA | SEQUENCE |
| --- | --- |
| mITGB3-1 | rCrUrArGrGrCrArArGrArArCrArUrUrArCrCrArArCrUrGAT |
| mITGB3-2 | rArGrCrUrGrArCrGrGrArUrArCrUrGrGrCrArArArArArCGC |
| circRAB12-1 | rArUrArUrArUrGrCrArArUrCrCrUrGrGrUrGrUrUrGrArCTT |
| circRAB12-2 | rArUrCrCrUrGrGrUrGrUrUrGrArCrUrUrCrArArArArUrCAA |
| circBACE2-1 | rGrArGrCrCrCrCrUrGrUrGrCrArGrUrArCrArGrArUrUrCTC |
| circBACE2-2 | rCrGrArGrCrCrCrCrUrGrUrGrCrArGrUrArCrArGrArUrUCT |
| circBIRC62-1 | rCrUrGrArUrGrArArCrCrUrUrGrUrArArArCrCrArGrGrUGG |
| circBIRC62-2 | rUrGrArUrGrArArCrCrUrUrGrUrArArArCrCrArGrGrUrGGA |

